# Vsb1, Ypq1 and Ypq2 control dynamic cationic amino acid storage in the yeast vacuole

**DOI:** 10.1101/2025.09.11.675521

**Authors:** Evi Zaremba, Fabienne Vierendeels, Raphaël Dutoit, Elisabeth Bodo, Ersilia Bifulco, Catherine Tricot, Elad Noor, Bruno André, Evgeny Onischenko, Melody Cools

## Abstract

Although the yeast vacuole plays a crucial role in storing and mobilizing cationic amino acids (CAA), CAA transport at the vacuolar membrane remains poorly understood. Here, by combining analysis of CAA pools, uptake and permeabilization assays, we establish Vsb1 as the principal vacuolar lysine transporter, enabling its strong accumulation in the vacuole while mitigating its toxicity. We further show that, although Ypq1 can mediate proton-independent vacuolar lysine import, it mainly functions as a lysine exporter necessary for lysine mobilization under conditions of lysine scarcity and downregulated as lysine stores are exhausted. Using quantitative models based on dynamic metabolic labeling, we further show that, surprisingly, in growing cells, CAA rapidly exchange between vacuolar and cytosolic compartments, a process involving the export activity of Ypq1 and its paralogue Ypq2, specific for lysine and arginine, respectively. Together, our findings reveal the unexpectedly complex function of Vsb1 and Ypq1/2 as the key transporters mediating dynamic vacuolar CAA storage.

**40-word summary:** Zaremba *et al*. characterize Vsb1 as the main yeast vacuolar lysine importer and Ypq1/2 as bi- directional vacuolar transporters of lysine and arginine. Their study highlights the role of vacuolar transporters in regulating cationic amino acid homeostasis under fluctuating nutrient availability.

## Introduction

Microorganisms inhabit dynamic environments and face numerous nutritional challenges. In nutrient-poor conditions, they must contend with intense competition for essential resources such as nitrogen, phosphorus, and trace metals (Broach, 2012; Sun et al., 2022; Watanabe et al., 2008). This requires highly efficient scavenging mechanisms (Kim and Klionsky, 2000), specific transport systems (Horák, 1997; Kschischo et al., 2016) and the capacity to store large quantities of metabolites (Klionsky, 1990) when nutrients are abundant as well as to recycle building blocks of macromolecules during periods of scarcity (Li and Vierstra, 2012; Reggiori and Klionsky, 2013). However, these processes require precise regulation when conditions shift to nutrient-rich environments, as excessive metabolite accumulation can result in toxicity (Li and Kane, 2009).

In budding yeast (*Saccharomyces cerevisiae*), the vacuole serves as a principal storage organelle for a variety of metabolites, including amino acids, polyphosphates, and metal ions (Klionsky, 1990), allowing cells to withstand long periods of nutritional deprivation (Li and Kane, 2009). Furthermore, because it can sequester harmful compounds, the vacuole significantly contributes to detoxification and resistance to toxic metabolites (Cools et al., 2020; Jézégou et al., 2012; Li and Kane, 2009). Functionally analogous to mammalian lysosomes, the yeast vacuole also hosts hydrolases that degrade macromolecules (Klionsky, 1990; Li and Kane, 2009) and sometimes entire organelles delivered via autophagy (Reggiori and Klionsky, 2013). Its dual role in storage and metabolite recycling has presumably driven the evolution of specialized vacuolar transport systems (Li and Kane, 2009).

The activity of most vacuolar importers is thought to depend on the proton gradient generated by the V-ATPase (Cools et al., 2020; Kawano-Kawada et al., 2019, 2021; Sekito et al., 2014b; Shimazu et al., 2005), a multiprotein complex that hydrolyzes ATP to ADP to pump protons into the vacuole. Its Vph1 subunit is essential for both assembly and activity (Li and Kane, 2009; Nishi et al., 2003). Among vacuolar transporters, arginine transporters have been extensively studied (Sekito et al., 2014a; Shimazu et al., 2005). Arginine import is primarily mediated by Vsb1, a member of the SLC26/SulP family of transporters (Cools et al., 2020). Unlike Vsb1, which transports positively charged arginine, most of the characterized transporters of the SLC26A/SulP family transport inorganic anions (Alper and Sharma, 2013). Members of this family are characterized by a transporter domain of 14 membrane-spanning ⍺-helices (TM) (Chi et al., 2020; Walter et al., 2019) and a cytosolic Sulfate Transport Anti-Sigma antagonist (STAS) domain (Chi et al., 2020; Sharma et al., 2011; Walter et al., 2019). Multiple lines of evidence suggest that these transporters act as homodimers (Bertoni et al., 2017; Chi et al., 2020; Walter et al., 2019; Waterhouse et al., 2018). Interestingly, Vsb1 features a long cytosolic N-terminal region and an additional RmlC-like fold domain of unknown function adjacent to its STAS domain (Giraud et al., 2000).

During steady state growth in a rich nitrogen medium, arginine is imported into the vacuole by Vsb1 and stored at high concentrations, which apparently requires negatively charged polyphosphate chains that are synthesized by the vacuolar transporter chaperone (VTC) complex (Cools et al., 2020; Dürr et al., 1979; Hothorn et al., 2009). However, under nitrogen starvation, Vsb1 activity is inhibited and Ypq2 exports arginine from the vacuole to the cytosol, where it serves as an alternate nitrogen source, crucial for long-term survival (Cools et al., 2020). Ypq2 belongs to the PQ-loop family of transporters, which are characterized by a conserved repeated PQ motif critical to their activity. Its human ortholog, PQLC2, is a lysosomal transporter that exports CAA (Jézégou et al., 2012). The coordinated actions of Vsb1 and Ypq2 enable yeast to adapt readily to changes in nitrogen availability (Cools et al., 2020).

Similarly to arginine, lysine is also predominantly stored in the vacuole (Messenguy et al., 1980). Its uptake into the cell is mediated by the high-affinity plasma membrane permease Lyp1, as well as Can1 and Gap1 under specific conditions (Grenson, 1966; Grenson et al., 1966; Sychrova et al., 1993). Lysine biosynthesis is tightly regulated: the first step, catalyzed by the homocitrate synthases encoded by *LYS20* and *LYS21*, combines ⍺-ketoglutarate and acetyl-CoA to form homocitrate and is subject to feedback inhibition by lysine (Feller et al., 1999; Ramos et al., 1996). In parallel, the transcription factor Lys14 induces the expression of lysine biosynthetic genes in response to lysine deprivation. Among Lys14 target genes is *LYS9* which encodes saccharopine dehydrogenase, an enzyme that catalyzes the seventh step of the pathway (Feller et al., 1999; Ljungdahl and Daignan-Fornier, 2012; Ramos et al., 1988). Deletion of any gene in this pathway renders cells lysine auxotrophs, dependent on external lysine for growth (Sinha and Bhattacharjee, 1971). Interestingly, unlike most amino acids, lysine cannot serve as a nitrogen source for yeast (Watson, 1976), although it can act as a precursor for high- level synthesis of polyamines (Olin-Sandoval et al., 2019). In addition, lysine can be toxic to yeast in certain contexts, such as growth on poor nitrogen sources although the reasons for this toxicity remain unclear (Cooper et al., 1979; Sumrada and Cooper, 1978; Thomas and Ingledew, 1994).

Various approaches have been employed to investigate lysine transport across the vacuolar membrane. In the 1970s and 1980s, researchers developed protocols to isolate intact vacuoles and vacuolar vesicles, enabling *in vitro* measurements of radiolabeled CAA uptake (Ohsumi et al., 1988; Wiemken and Nurse, 1973). These studies, particularly those exploiting isolated vacuoles derived from mutant strains, provided genetic evidence that Vba1, Vba2, and Vba3, as well as Ypq1 and Ypq3, two transporters closely related to Ypq2, are involved in CAA transport at the vacuolar membrane(Jézégou et al., 2012; Kawano-Kawada et al., 2019; Manabe et al., 2016; Sekito et al., 2014b; Shimazu et al., 2005). Given that 80 to 90% of the total soluble CAA pool is stored in the vacuole, total pool measurements have also been widely used to investigate the impact of mutations on vacuolar lysine accumulation (Kitamoto et al., 1988; Messenguy et al., 1980). Studies based on such analyses have suggested roles for Vsb1 in the vacuolar transport of lysine and histidine in addition to arginine (Kawano-Kawada et al., 2021), and a role of Avt4 in lysine export from the vacuole (Sekito et al., 2014a). Additional insights have been provided by permeabilization-based assays, such as the cupric ion and cytochrome C methods, which allow differential extraction of intracellular amino acid pools (Ohsumi et al., 1988; Wiemken and Nurse, 1973). Most recently, Arines *et al*. developed a protocol for reconstituting purified vacuolar transporters into liposomes, enabling the direct detection of Ypq1-mediated ^14^C-lysine transport (Arines et al., 2024). Thus, even if several candidate proteins have been proposed to mediate lysine transport at the vacuolar membrane, direct physiological evidence identifying the primary lysine importer and exporter *in vivo* remains limited. Moreover, how vacuolar CAA transporters work together to maintain cellular CAA homeostasis under different physiological conditions is not well understood.

In this study we combine structural modeling coupled with mutagenesis experiments, biochemical transport assays and quantitative models based on dynamic labeling in live cells to explore the dynamics of vacuolar CAA transport during active growth and under stress conditions. We demonstrate that, under steady state conditions, in addition to its established role in arginine transport, Vsb1 acts as the main proton-gradient dependent importer responsible for vacuolar accumulation of lysine. Its RmlC-like fold domain and key residues Asp-223, Tyr-227 and Glu-278 within the transmembrane domain were found to be essential for its activity. In contrast to import, vacuolar export of lysine and arginine follow separate routes, preferentially mediated respectively by Ypq1 and Ypq2, which under steady-state growth conditions enable surprisingly high export rates corresponding to approximatively one total pool of cellular CAA per hour. Additionally, we show that while Vsb1 accumulates lysine in the vacuole to shield cells from the toxic effects of its excess, vacuolar export mediated by Ypq1 is essential for mobilizing vacuolar lysine stockpile during its scarcity.

## Results

### Vsb1 and Ypq1 are involved in lysine transport across the vacuolar membrane

We and others have previously demonstrated that arginine transport at the vacuolar membrane is catalyzed by the putative proton antiporter Vsb1 and by the Ypq2 exporter (Cools et al., 2020; Kawano-Kawada et al., 2021). However, the mechanisms of vacuolar transport of lysine, another major CAA, remain only partially characterized. As most candidate vacuolar lysine transporters have been proposed to function as proton antiporters (Cools et al., 2020; Sato et al., 1984; Sekito et al., 2014b), we investigated whether lysine accumulation in the vacuole requires the vacuolar proton gradient (Sato et al., 1984). Thus, we analyzed the effect of the *VPH1* deletion on the total pool of soluble lysine (Li and Kane, 2009; Manolson et al., 1992; Preston et al., 1989). As shown by previous studies, this analysis reliably reflects vacuolar lysine content, as at least 80–90% of free cellular lysine resides in the vacuole (Cools et al., 2020; Messenguy et al., 1980). UPLC analysis of soluble metabolites revealed that deletion of *VPH1* resulted in a significant reduction in soluble lysine content (Fig. 1), implying that lysine accumulation requires the proton gradient and that vacuolar lysine importers are likely proton dependent.

**Figure 1.**
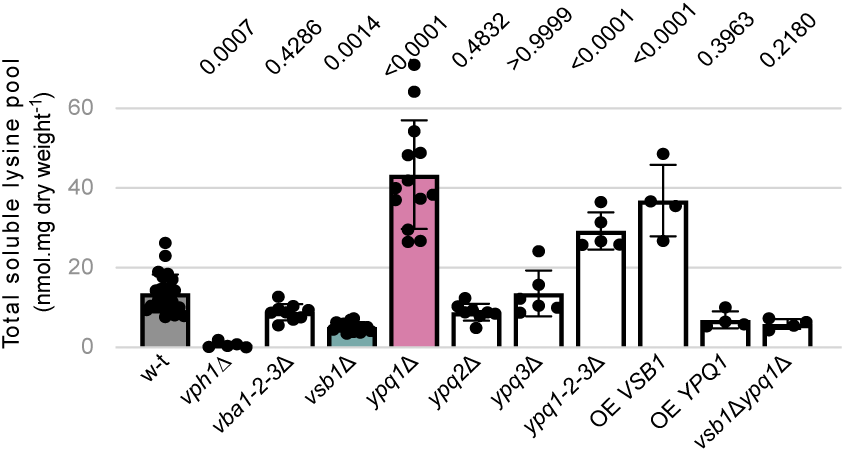
Vsb1 and Ypq1 are involved in lysine transport across the vacuolar membrane The intracellular lysine content was measured in the w-t, *vph1*Δ, *vba1-2-3*Δ, *vsb1*Δ, *ypq1*Δ, *ypq2*Δ, *ypq3*Δ, *ypq1-2-3*Δ, OE VSB1 (*vsb1*Δ complemented with FV438 plasmid, allowing *VSB1* overexpression under TDH3 promoter), OE YPQ1 (ypq1Δ complemented with FV445 plasmid, allowing YPQ1 overexpression under TDH3 promoter) and *vsb1*Δ*ypq1*Δ strains. The p-values were calculated by a one-way ANOVA with post-hoc comparison tests with the w-t strain (n = 4-24).

We next sought to investigate the role of eight putative lysine transporters in vacuolar lysine accumulation: Avt4 (Russnak et al., 2001; Sato et al., 1984; Sekito et al., 2014a; Yang et al., 2006), Vba1-2-3 (Ohsumi and Anraku, 1981; Sato et al., 1984; Shimazu et al., 2005), Vsb1 (Cools et al., 2020; Kawano-Kawada et al., 2021) and Ypq1-2-3 (Arines et al., 2023; Jézégou et al., 2012; Manabe et al., 2016; Sekito et al., 2014b). Among them, Avt4 was discarded because our reference strain (Σ1278b) carries an insertion that leads to a premature stop codon in the *AVT4* gene (Cherry et al., 2012). We therefore focused on the remaining seven proteins and assessed their contribution to lysine accumulation by measuring the total soluble lysine pool in the corresponding deletion mutants (Fig. 1). Among the tested mutants, the *vsb1Δ* strain displayed the strongest reduction in the intracellular lysine pool, pointing to a defect in the vacuolar accumulation of lysine. Interestingly, the *ypq1Δ* and triple *ypq1-2-3Δ* mutants showed the opposite effect, namely an over-accumulation of intracellular lysine, which may point to an impaired lysine mobilization from the vacuole. The other analyzed deletion mutants (*ypq2Δ*, *ypq3Δ*, *vba1-2-3Δ*) displayed close to no impact on lysine accumulation, meaning that Vsb1 and Ypq1 are likely the key players in lysine transport across the vacuolar membrane.

To further assess the role of Vsb1 and Ypq1 in the establishment of cellular lysine levels, total soluble lysine pool measurements were additionally performed in strains transformed with a plasmid expressing *VSB1* or *YPQ1* under the strong constitutive *TDH3* promoter (McAlister and Holland, 1985). *VSB1* overexpression led to a 3-fold increase in the total lysine pool, whereas *YPQ1* overexpression reduced it to levels comparable to that of the *vsb1Δ* strain (Fig. 1). These findings further support the conclusion that Vsb1 and Ypq1 play important roles in vacuolar lysine transport.

We finally examined the epistasis relationship between *vsb1Δ* and *ypq1Δ* deletions by comparing the total lysine pool of the double *vsb1Δypq1Δ* mutant to those of the previously analyzed single mutant strains. The double mutant displayed similar soluble lysine levels as the *vsb1Δ* mutant (Fig. 1), indicating that the effect of *VSB1* loss is epistatic over that caused by the *YPQ1* deletion. Thus, Vsb1 likely acts upstream of Ypq1 in the control of vacuolar lysine levels.

### Vsb1 mediates proton gradient dependent vacuolar accumulation of lysine

Next, we analyzed in more detail the role of Vsb1 in the import and accumulation of lysine in the vacuole. To achieve this, we first measured the total free lysine content in the wild type (w-t) and the *vsb1*Δ strains before and after addition of 500 µM lysine. Although exogenous lysine accumulated intracellularly in both cases, the *vsb1*Δ cells accumulated lysine much slowly and to lower total concentrations (8.5-fold less) as compared to the w-t strain (Fig. 2 A). A plausible explanation for this impairment is an excessive accumulation of lysine in the cytosol of the *vsb1*Δ mutant, which in turn downregulates plasma membrane transporters (see below).

**Figure 2.**
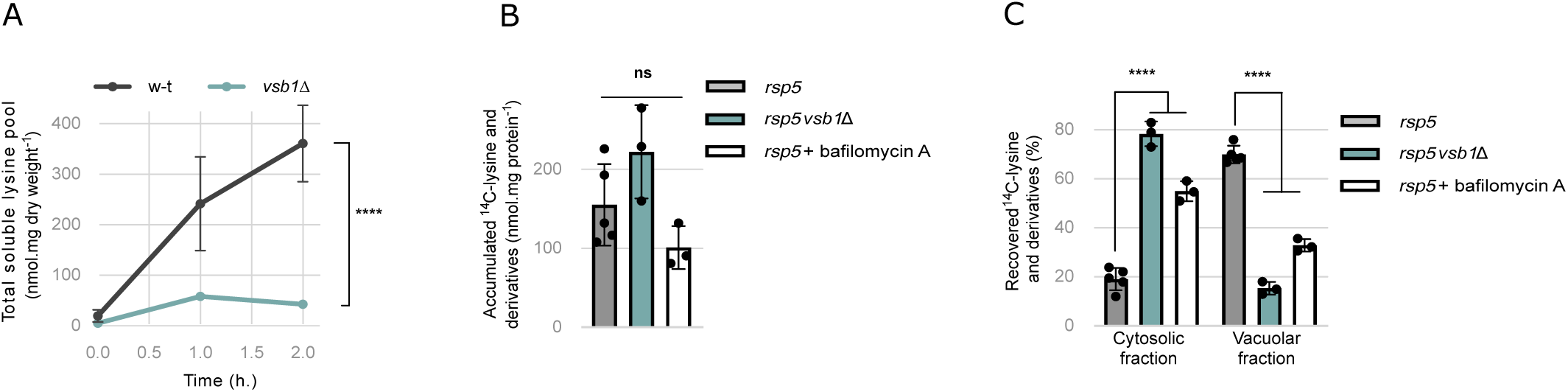
Vsb1 mediates proton gradient dependent vacuolar accumulation of lysine. (A) The intracellular lysine content was measured in the w-t and *vsb1*Δ strains after addition of 500 µM lysine in the media. (****: p < 0.0001 by two-way ANOVA test) (n = 4). (B) The *^14^*C-lysine accumulated in the *rsp5* and *rsp5vsb1*Δ strains (30 μM and 90 μM *^14^*C-lysine, respectively) and the *rsp5* strain treated with bafilomycin A (9 μM) for 15 min prior to the assay (90 μM *^14^*C-lysine) (ns: p > 0.05 by one-way ANOVA test) (n = 3-5). (C) Distribution of initially accumulated *^14^*C-lysine and derivatives between the cytosolic and vacuolar fractions after cell permeabilization with cytochrome C. (****: p < 0.0001 by one-way ANOVA test) (n = 3-5).

To test whether lysine accumulates in the cytosol in the absence of Vsb1, we incubated cells with ^14^C-labeled lysine, and then selectively permeabilized the plasma membrane using cytochrome C in order to successively extract cytosolic and vacuolar fractions (Cools et al., 2020; Messenguy et al., 1980; Wiemken and Nurse, 1973). For these experiments we used a hypomorphic *rsp5* mutant background, in which plasma membrane transporters accumulate as a consequence of reduced expression of the Rsp5 ubiquitin ligase (Hein et al., 1995). To allow for direct comparison, the concentrations of ^14^C-lysine added to cells were adjusted, ensuring similar total lysine uptake (Fig. 2 B). In the *rsp5* mutant, over 75% of ^14^C-lysine and its derivatives were vacuolar. In contrast, in *rsp5vsb1Δ* cells the vacuolar content of ^14^C-lysine dropped to 20% while approximately 80 % was retained in the cytosol. This confirms that Vsb1 contributes to the vacuolar import of exogenous lysine in growing cells (Fig. 2 C). To show Vsb1-dependent sequestration of lysine is dependent on the proton gradient, *rsp5* cells were treated with V- ATPase inhibitor bafilomycin A prior to the addition of ^14^C-lysine and permeabilization. We again adjusted exogenous lysine concentrations to reach comparable intracellular accumulation (Fig. 2 B). Treating cells with bafilomycin A resulted in a pronounced shift of ^14^C-lysine and its derivatives towards the cytosolic fraction (Fig. 2 C), in line with the hypothesis that Vsb1- dependent lysine accumulation into the vacuole requires the proton gradient established by the V-ATPase.

### Vacuolar accumulation of lysine mediated by Vsb1 attenuates Lyp1 downregulation and mitigates lysine toxicity

To further investigate the connection between defect of vacuolar accumulation of lysine in the *vsb1Δ* mutant and its reduced cellular uptake (Fig. 2 A), we measured the effect of bafilomycin A and *VSB1* deletion on the kinetics of ^14^C-lysine accumulation. The initial uptake rates were equivalent across all conditions (Fig. 3 A), indicating that the plasma membrane lysine transporter, Lyp1 (Jauniaux and Grenson, 1990; Sychrova et al., 1993), remained comparably active. However, long-term accumulation of ^14^C-lysine was strongly reduced in the *vsb1*Δ strain or upon bafilomycin A treatment (Fig. 3 B). This confirms that external lysine accumulation seems connected to its vacuolar import.

**Figure 3.**
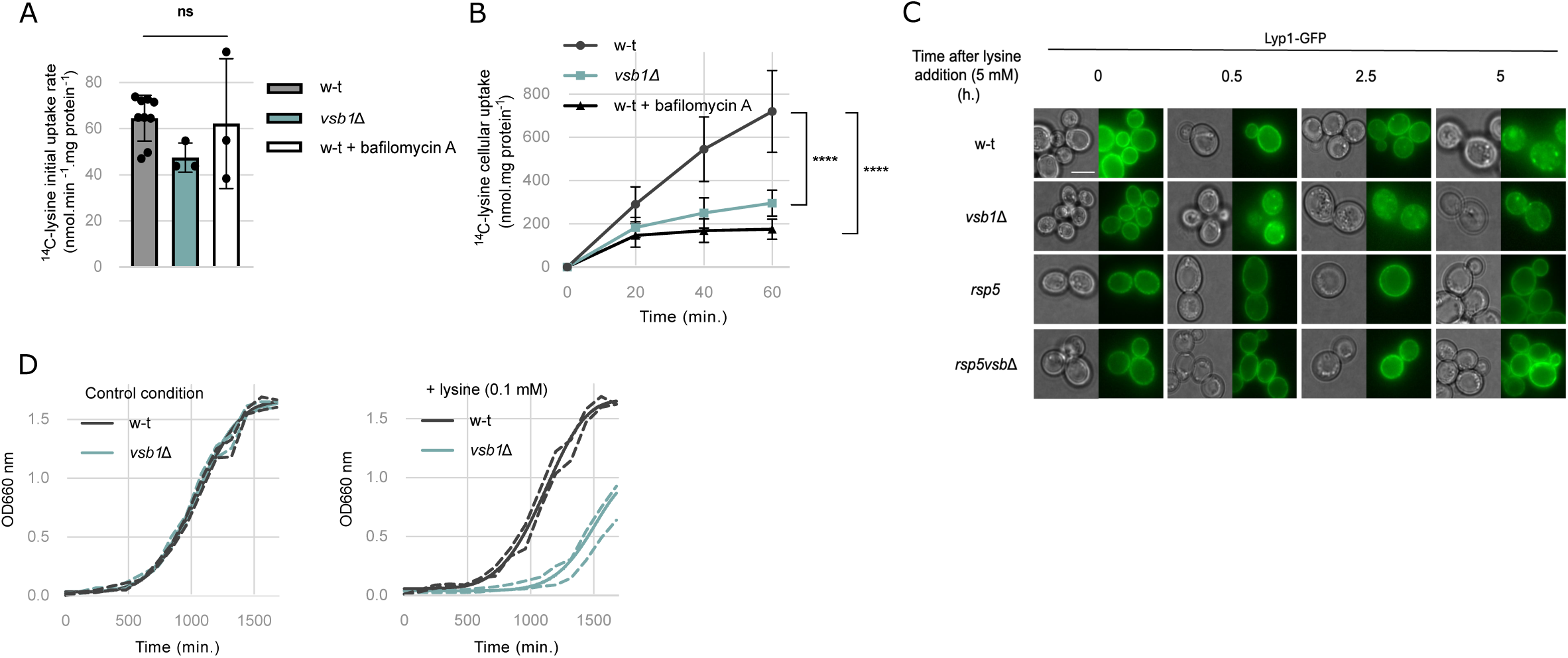
Vacuolar accumulation of lysine mediated by Vsb1 attenuates Lyp1 downregulation and mitigates lysine toxicity. (A) Initial uptake rate of ^14^C-lysine (500 µM) in w-t and *vsb1*Δ cells and w-t cells treated with bafilomycin A (9 mM) for 15 min prior to the assay (ns: p > 0.05 by one-way ANOVA) (n = 3-9). Initial uptake rates were determined during the first 90 s interval after ^14^C-lysine addition. (B) Time course accumulation of ^14^C-lysine (500 µM) over 60 min into w-t and *vsb1*Δ cells and w-t cells treated with bafilomycin A (9 mM) for 15 min before the uptake assay (****: p < 0.0001 by two-way ANOVA) (n = 3-9). (C) Microscopy analysis of Lyp1-GFP expressed from a plasmid in w-t, *vsb1*Δ, *rsp5* and *rsp5vsb1*Δ strains before and after 500 µM lysine addition in the culture media (0.5, 2.5 and 5 h). Scale: 5 µm. (D) Growth of the w-t and *vsb1*Δ strains in the absence or presence of lysine (100 µM) in the culture media. Cells were initially grown on minimal lysine-free medium containing proline (10 mM) as the sole nitrogen source and the optical density at 660 nm (OD 660 nm) was monitored over a 24-hour period. Growth is represented as the Weibull nonlinear fit curve with the 99 % confidence bands (n = 3).

To assess whether reduced lysine accumulation in *vsb1*Δ cells was due to the downregulation of Lyp1, we expressed a C-terminally GFP-tagged Lyp1 from a plasmid, in w-t and *vsb1*Δ strains. Lyp1-GFP localized predominantly at the plasma membrane in both strains in the absence of lysine (Fig. 3 C). However, upon the addition of external lysine, Lyp1-GFP was internalized, and this occurred significantly faster in the *vsb1*Δ mutant than in the w-t cells (Fig. 3 C). In *rsp5* mutant backgrounds, where endocytosis of plasma membrane transporters is impaired (Hein et al., 1995), this internalization was blocked, confirming that it occurs via endocytosis. Considering our observations that the *vsb1*Δ mutant accumulates exogenous lysine in the cytosol (Fig. 2 C) these results support a model in which defective lysine import into the vacuole, as in the *vsb1*Δ mutant, leads to an excessive accumulation of lysine in the cytosol that subsequently triggers an early endocytosis of Lyp1.

As lysine has been previously shown to be toxic for yeast cells grown on a poor nitrogen source (Sumrada, 1976), we investigated how Vsb1-mediated lysine compartmentalization in the vacuole could contribute to protecting cells from lysine toxicity. To this end, we monitored the growth of w-t and *vsb1*Δ strains in proline medium with or without lysine supplementation. Supplementation with 100 µM lysine did not affect the growth of the w-t strain (Fig. 3 D). In contrast, the *vsb1*Δ mutant exhibited a longer lag phase as well as a reduced growth rate when grown in the presence of lysine. When ammonium was used as a nitrogen source, the hypersensitivity of the *vsb1*Δ strain lysine was less pronounced (Fig. S 1), likely because the Gap1 permease involved in Lys uptake is less active under these conditions (Grenson, 1983; Merhi and André, 2012). These observations support the importance of vacuolar accumulation of lysine mediated by Vsb1 in maintaining lysine homeostasis and protecting cells from its toxic effects.

### The cytosolic RmlC-like domain of Vsb1 is essential for its CAA transport activity

To gain further insight into the mechanism of Vsb1 function, we set out to investigate the structural basis of its CAA transport function. Evolutionarily, Vsb1 belongs to the SLC26A/SulP superfamily of anion exchangers or anion channels ubiquitous in all kingdoms of life (Alper and Sharma, 2013). According to several available structures, SLC26A/SulP transporters contain a transmembrane domain having 14 ⍺-helices and a STAS domain required for dimerization (Bavi et al., 2021; Butan et al., 2022; Chi et al., 2020; Futamata et al., 2022; Ge et al., 2021; Hu et al., 2024; Liu et al., 2023; Tippett et al., 2023; Walter et al., 2019; Wang et al., 2019, 2024, 2021). In the case of Vsb1, it has a cytosolic N-terminal tail and a putative RmlC-like domain in addition to the SLC26A/SulP-like transmembrane domain and the STAS domain (Fig. 4 A). Since the structure of Vsb1 has not yet been solved, we modeled its molecular architecture as a dimer using the AlphaFold 3 Server (Abramson et al., 2024). The full-length homodimer has a predicted template modeling score of 0.59, indicating overall poor structure prediction (Fig. S 2 A). However, several regions of the protein appeared disordered with a predicted confidence score below 0.5 (residues 1-188 and 818-848), indicating that either these regions are natively unfolded or fail to be properly modeled due to the lack of structural homologs in the Protein Data

**Figure 4.**
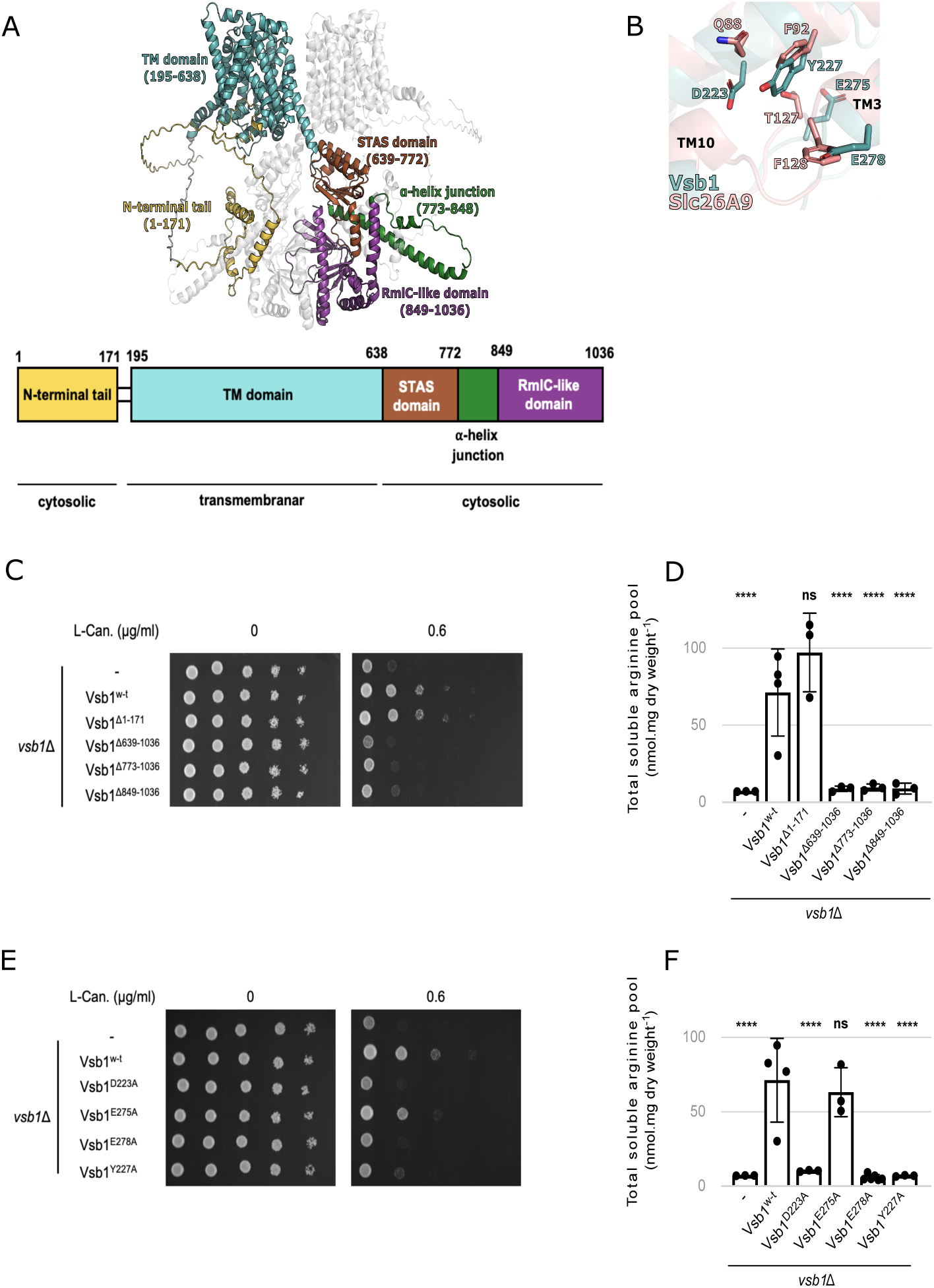
Insights into Vsb1 function from structural modeling and mutational analyses. (A) Overall representation of Vsb1 structure modeled as a dimer using AlphaFold 3. The domains and regions of monomer A are represented as: N-terminal tail in yellow, the transmembrane (TM) domain in teal, the STAS domain in red, the α-helix junction in green, and the RmlC-like domain in purple. Monomer B is represented in light grey. (B) Close-up view of the putative CAA binding site of Vsb1 (teal) and comparison with the Cl- binding site of the mammalian Slc26A9 (pink). The residues involved in chloride transport in Slc26A9 (Walter et al., 2019) and their Vsb1 counterparts are annotated and shown in stick representation. The R.M.SD. of the structural alignment Vsb1 model with Slc26A9 (PDB code 6RTC) is 3.76 Å. (C) The effect of Vsb1 truncations was evaluated by assessing canavanine susceptibility of a *vsb1*Δ strain transformed with an empty plasmid (-) or a plasmid allowing the expression of different *VSB1* truncation mutants under the *VSB1* native promoter. Cells were spread on minimal medium containing 0.6 µg/ml canavanine or not and cell growth was evaluated after a 3-days incubation at 29°C. (D) The intracellular arginine content was measured for the *vsb1*Δ strains expressing the different *VSB1* truncations (ns: p > 0.05; ****: p < 0.0001 by one-way ANOVA with post-hoc comparison tests with the w-t strain) (n = 3-4). (E and F) A *vsb1*Δ strain was transformed with an empty plasmid (-) or plasmids expressing different *VSB1* variants under the *VSB1* native promoter. Cells spread on minimal medium containing canavanine or not were incubated for 3 days at 29°C (E) and their intracellular arginine content was measured (F). (ns: p > 0.05; ****: p < 0.0001 by one-way ANOVA with post-hoc comparison tests with the w-t strain) (n = 3-4).

Base (Terwilliger et al., 2024). Indeed, when these poorly modeled regions are omitted, the Vsb1 model exhibited a much better confidence score, between 0.7 and 0.9, compatible with its functional interpretation (Fig. S 2 A).

As expected, the predicted folds of the transmembrane domain (195-638) and STAS domain (639-772) are closely related to the established SLC26A/SulP transporter structures. The former is composed of 14 transmembrane ⍺-helices (TM) segregated into a gate and a core subdomain (Fig. S 2 B). The putative substrate binding pocket is lodged between TM3 and TM10 (Fig. 4 B and Fig. S 2 B). The predicted interaction between the monomers in the dimer model is mediated partially by the STAS domains with an interaction surface of 385 Å^2^. Since the STAS domains are swapped between monomers, the STAS domain of a monomer seems to also interact with the transmembrane domain of the other monomer (Fig. 4 A). Finally, the fourth domain of Vsb1 (773- 1036) is predicted to adopt an RmlC-like jelly roll fold (849-1036) and to connect to the STAS domain by a long junctional ⍺-helix (774-848). Such a fold is often associated with proteins that bind cyclic nucleotide, nucleotide-activated sugar, or metal ion (Dunwell et al., 2001). The RmlC- like domain can also contribute to protein multimerization as observed for the mammalian potassium channel Eag1 (Whicher and MacKinnon, 2016). In Vsb1, it could possibly play a role in dimerization since it provides a massive projected interaction surface between monomers of 2180 Å^2^. The presence of such a domain in a transporter has only been reported for LtnT, a SLC26A/SulP transporter involved in nitrate transport from the cyanobacterium *Synechococcus elongatus* (Maeda et al., 2006). It is worth noticing that Vsb1 lacks the intervening sequence and the PDZ domain found in mammalian SLC26A/SulP transporters.

To investigate the functional significance of the cytosolic domains, we generated Vsb1- truncation variants that encompass the N-terminal tail (Vsb1^Δ1-171^), the RmlC-like domain (Vsb1^Δ849-1036^), the RmlC-like domain with the ⍺-helix junction (Vsb1^Δ773-1036^) or the entirety of the C-terminal region (Vsb1^Δ639-1036^) (Fig. 4 A). First, we analyzed their intracellular localization using the C-terminal GFP fusions expressed under the *VSB1* gene’s native promoter in a *vsb1Δ* deletion background. The truncation variants were expressed mainly as full-length proteins although at somewhat different levels (Fig. S 3 B). Most of them were also efficiently targeted to the vacuolar membrane as shown by colocalization with lipophilic dye FM4-64 used as a fiducial vacuolar membrane marker (Vida and Emr, 1995) (Fig. S 3 A). Next, to test the functionality of the truncated variants we expressed them in an untagged form in the *vsb1Δ* deletion background as GFP fusion was shown to be incompatible with Vsb1 activity (Cools et al., 2020; Kawano-Kawada et al., 2021) and tested them for supporting growth on plates containing canavanine, an analogue of arginine to which yeast cells lacking *VSB1* are hypersensitive (Cools et al., 2020) (Fig. 4 C). Whereas complementing *vsb1Δ* mutant with Vsb1^Δ1-171^ supported cell growth in the presence of canavanine similarly to w-t Vsb1, the other truncation variants failed to complement the *vsb1Δ* mutant. Consistent with these observations, the same truncations impaired the intracellular accumulation of CAA (Fig. 4 D and Fig. S 3 D-E). Together these results indicate that the cytosolic domains are essential for Vsb1 activity and a critical part can be mapped to the RmlC-like domain (residues 849-1036) (Fig. 4 C).

### Asp-223, Tyr-227 and Glu-278 of the putative CAA binding site are essential for Vsb1 activity

Next, we sought to investigate the requirement of specific residues within the transmembrane domain of Vsb1 for CAA transport, explicitly in light of the fact that members of the SLC26A/SulP superfamily are classically described as anion transporters. In mammalian SLC26A transporters, the anion binding site is globally positively charged via the macro-dipoles of TM3 and TM10 (Futamata et al., 2022; Ge et al., 2021; Hu et al., 2024; Liu et al., 2023; Tippett et al., 2023; Wang et al., 2021) and within this site an arginine residue plays a crucial role in anion binding for SLC26A3, SLC26A4, SLC26A5, and SLC26A6 (Futamata et al., 2022; Gorbunov et al., 2014; Liu et al., 2023; Tippett et al., 2023; Wang et al., 2024, 2021). Therefore, there must be some adaptations in Vsb1 to allow the binding of CAA. A previous study has identified Asp-223, which is conserved among the fungal Vsb1 orthologues, as being necessary for transport of CAA (Kawano-Kawada et al., 2021; Ohnishi et al., 2022). Based on our model, Asp-223 is conveniently located in the putative binding site between TM3 and TM10 (Fig. 4 B). In the characterized mammalian transporters of SLC26A/SulP family, a glutamine residue is found at a position equivalent to Asp-223 and implicated in anion interaction (Fig. 4 B and Fig. S 2 C) (Futamata et al., 2022; Ge et al., 2021; Liu et al., 2023; Tippett et al., 2023; Walter et al., 2019; Wang et al., 2024). In SLC26Dg from the bacterium *Deinococcus geothermalis,* a glutamate residue equivalent to Asp-223 is required for the fumarate/Na^+^ exchange (Geertsma et al., 2015). By comparing the Vsb1 model with the related SLC26A transporters, we identified three other candidate residues that could be involved in CAA transport: Tyr-227, Glu-275 and Glu-278 (Fig. 4 B and Fig. S 2 C). Of note, the arginine residue found in the anion binding site of some SLC26A/SulP transporters is absent in Vsb1 (Fig. S 2 C). In all structurally characterized SLC26A members, an aromatic residue is found within the substrate binding site at a position equivalent to Tyr-227 and plays a role in anion exchange (Butan et al., 2022; Futamata et al., 2022; Hu et al., 2024; Terwilliger et al., 2024; Tippett et al., 2023). Glu-275 is positioned in the second shell of interaction, close to Tyr-227, possibly imposing steric constraint on the tyrosyl group. Finally, Glu-278 is localized at the N-terminal end of TM3 and could neutralize the helix macro-dipole. It has been proposed that the helix macro-dipoles of TM3 and TM10 play a role in anion binding in some SLC26A transporters (Geertsma et al., 2015; Terwilliger et al., 2024).

To investigate the roles of these four candidate key residues, Vsb1 variants were constructed, each of them separately substituted with an alanine residue. As in the case of truncations, the GFP-fused mutants correctly localized to the vacuolar membrane and were expressed at similar levels to the w-t variant (Fig. S 3 A and 3 C). The untagged variants were then tested for their ability to rescue the canavanine hypersensitivity (Fig. 4 E) and reduced levels of CAA of the *vsb1Δ* deletion mutant (Fig. 4 F and Fig. S 2 D-E). While Vsb1^E275A^ variant was only slightly more sensitive to canavanine than the w-t and had levels of cellular CAA similar to the w-t cells, the Vsb1^D223A^, Vsb1^Y227A^ and Vsb1^E278A^ variants failed to rescue both phenotypes. These observations strongly support the critical role of Asp-223, Tyr-227 and Glu-278 in CAA transport by Vsb1.

### Ypq1 is essential for lysine mobilization from the vacuole under lysine starvation

Our observation that lysine accumulates to higher levels in the *ypq1*Δ mutant and that its overexpression reduced the lysine pool (Fig. 1) is consistent with Ypq1 acting as a vacuolar lysine exporter. To clarify the role of Ypq1 *in vivo*, we first assessed the activation of the lysine biosynthesis pathway in both w-t and *ypq1*Δ strains after lysine withdrawal, reasoning that this response could serve as an indicator of the availability of cytosolic lysine. To monitor this response, cells were grown in a lysine-supplemented medium and then washed and resuspended in a lysine-free medium all while monitoring mRNA levels of *LYS9*, the gene that encodes saccharopine dehydrogenase involved in lysine biosynthesis and that is known to be strongly induced upon lysine deprivation (Storts and Bhattacharjee, 1987). In the presence of lysine, the expression of *LYS9* was comparable in both w-t and *ypq1Δ* strains and, as expected, was induced after lysine withdrawal (Fig. 5 A). However, in the *ypq1*Δ mutant, this induction occurred earlier and was much more pronounced. This increased biosynthetic response is consistent with the view that lysine is released from the vacuole less efficiently in the absence of Ypq1.

**Figure 5.**
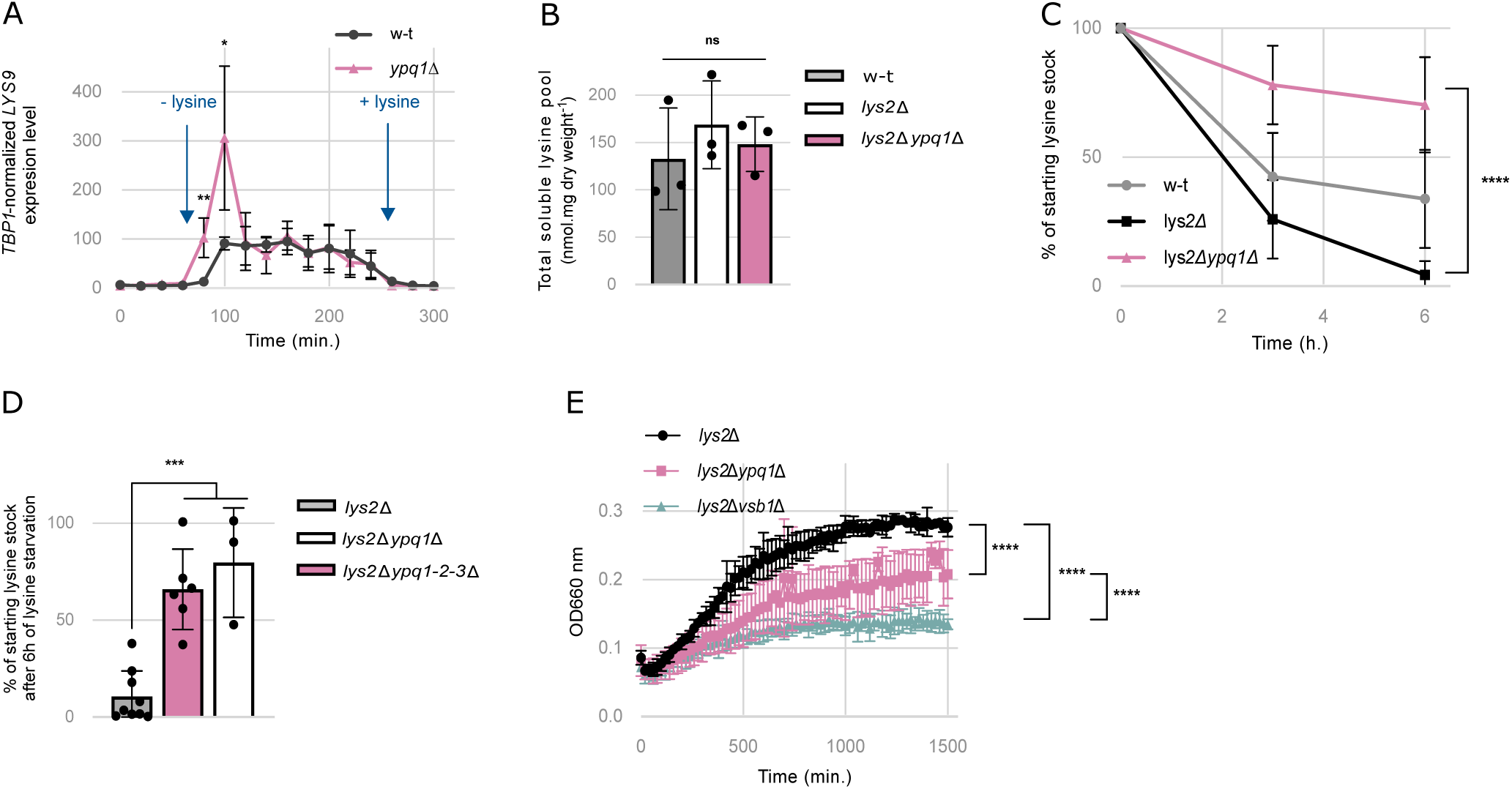
Ypq1 is essential for lysine mobilization from the vacuole under lysine starvation. (A) Impact of lysine on relative expression levels of *LYS9* in the w-t and *ypq1*Δ strains. Cells were grown in minimal medium supplemented with lysine. At the designated time points, cells were collected, washed and resuspended in a lysine-free minimal medium (-lysine), followed by re-supplementation with lysine (+lysine). Samples were collected every 20 min over a 5 h period for quantitative RT-PCR analysis of *LYS9* expression. (*: p < 0.05; **: p < 0.005 by Student’s t-test) (n = 4). (B) The intracellular lysine content was measured in the w-t, *lys2*Δ and *lys2*Δ*ypq1*Δ strains grown in minimal medium supplemented with lysine (ns: p > 0.05 by one-way ANOVA test) (n = 3). (C) Percentage of the initial intracellular lysine pool left after lysine starvation. The intracellular lysine content was measured in the w-t, *lys2*Δ and *lys2*Δ*ypq1*Δ strains 0, 3 and 6 h after transfer from a minimal medium supplemented with lysine to a lysine-free medium. (****: p < 0.0001 by one-way ANOVA test) (n = 4-12). (D) The intracellular lysine content was measured in the *lys2*Δ, *lys2*Δ*ypq1*Δ and *lys2*Δ*ypq1-2-3*Δ strains 0 and 6 h after transfer from a minimal medium supplemented with lysine to a lysine-free medium. (***: p < 0.001 by one-way ANOVA test) (n = 3-9). (E) Residual growth of the *lys2*Δ, *lys2*Δ*ypq1*Δ and *lys2*Δ*vsb1*Δ strains under lysine starvation. Cells were initially grown on minimal medium supplemented with lysine (416 µM) then collected by centrifugation and washed with lysine-free medium. Subsequently, cells were resuspended in lysine-free medium and the optical density (OD 660 nm) was monitored over a 24-hour period. (****: p < 0.0001 by two-way ANOVA test) (n = 3).

To test the role of Ypq1 in the vacuolar export of lysine more directly, we compared total soluble lysine pools of *lys2*Δ and *lys2*Δ*ypq1*Δ strains subjected to lysine starvation. Here the *LYS2* gene encoding ⍺-aminoadipate reductase was deleted in order to confer lysine auxotrophy (Feller et al., 1999; Sinha and Bhattacharjee, 1971). The w-t, *lys2*Δ and *lys2*Δ*ypq1*Δ strains were initially cultured in minimal medium supplemented with 500 µM lysine and then shifted to lysine-free medium. In the presence of external lysine, all three strains accumulated lysine at similar levels (Fig. 5 B). However, upon lysine withdrawal, total lysine pools rapidly declined in the w-t and the *lys2*Δ mutant strains but not in *lys2Δypq1Δ* double mutant which still retained approximately 65 % of its initial lysine content even after six hours of starvation (Fig. 5 C). We therefore conclude that lysine mobilization from the vacuole is impaired in the *lys2*Δ*ypq1*Δ mutant, indicating that Ypq1 mediates vacuolar lysine export under lysine scarcity.

Since the *lys2*Δ*ypq1*Δ mutant was still able to mobilize 35 % of its soluble lysine stores, other transporters could contribute to the export of lysine from the vacuole. Obvious candidates for this role include Ypq2 and Ypq3, two proteins evolutionarily related to Ypq1, which have been previously described as CAA transporters (Cools et al., 2020; Jézégou et al., 2012; Kawano- Kawada et al., 2019; Manabe et al., 2016). To determine whether Ypq2 or Ypq3 can mobilize lysine from the vacuole, we additionally analyzed the lysine stores of the *lys2*Δ*ypq1-2-3*Δ strain after 6 hours of lysine starvation. However, the quadruple mutant did not show any significant difference compared to the *lys2Δypq1Δ* strain (Fig. 5 D), indicating that neither Ypq2 nor Ypq3 effectively contribute to lysine mobilization in these conditions.

We next examined whether impaired export of the amino acid from the vacuole or defective accumulation prior to starvation affects the ability of the cell to sustain proliferation. We thus compared the proliferation capacity of *lys2Δ*, *lys2Δypq1Δ*, and *lys2Δvsb1Δ* strains following lysine withdrawal. The residual growth of the *lys2Δypq1Δ* mutant, which is defective in lysine mobilization, and the *lys2Δvsb1Δ* mutant, which stores lysine at much lower levels, was markedly reduced under lysine starvation compared to the *lys2Δ* strain (Fig. 5 E). This suggests that vacuolar lysine mobilization mediated by Ypq1 is an important means to maintain cell growth under lysine starvation.

Previous studies have reported vacuolar internalization and degradation of Ypq1 in response to lysine starvation (Arines et al., 2021; Li et al., 2015b). Considering our findings implicating Ypq1 in vacuolar lysine export during lysine starvation, we set out to investigate whether Ypq1 is degraded in our experimental conditions. To this end, Ypq1 was endogenously C-terminally tagged with GFP (Ypq1-GFP) in the w-t and the *lys2*Δ mutant. The Ypq1-GFP construct was functional as evidenced by the w-t levels of soluble intracellular lysine (Fig. S 4 A) and by the expected targeting of the GFP fusion to the vacuolar membrane observed in both strains grown in the presence of lysine (Fig. S 4 B and C). When cells were transferred to a lysine-free medium, Ypq1-GFP remained at the vacuolar membrane in the w-t strain, even after 6 hours. In contrast, three distinct cell populations were observed in the *lys2Δ* mutant: one in which Ypq1-GFP persisted at the vacuolar membrane, another in which it was targeted to the vacuolar lumen and a third in which an intermediate condition was observed (Fig. S 4 B and C). To verify that luminal GFP was associated with the degradation of Ypq1-GFP, cell extracts were collected under the same conditions and analyzed by western blotting (Fig. S 4 D and E). Following lysine withdrawal, Ypq1-GFP levels in the *lys2Δ* mutant showed a slight decrease, accompanied by an accumulation of free GFP (Fig. S 4 D and E). Notably, this did not occur in the lysine-prototrophic strain. These results confirm previous reports that Ypq1 is downregulated upon lysine starvation (Arines et al., 2021; Li et al., 2015b; Zhu et al., 2017), although this response appears to be partial and heterogeneous across the cell population. Interestingly, Ypq1 downregulation occurs hours after lysine withdrawal, when less than 20 % of the initial soluble lysine stock remains (Fig. 5 C), suggesting that Ypq1 downregulation occurs upon depletion of its intravacuolar substrate.

All in all, these results indicate that Ypq1 is the major vacuolar lysine exporter, necessary to mobilize the Vsb1-established lysine stores under lysine starvation conditions, dynamically regulated in correlation to vacuolar lysine availability.

### Ypq1 and Ypq2 mediate fast and semi-selective export of vacuolar lysine and arginine in actively growing cells

The fact that deletion of *YPQ1* and *YPQ2* leads to overaccumulation of lysine (Fig. 1) and arginine (Cools et al., 2020) respectively, in cells growing in replete medium, suggests that Ypq1 and Ypq2 can export CAA not only under starvation conditions but also during active cell growth. Additionally, given that both proteins are homologous to the mammalian PQLC2 transporter, which transports both lysine and arginine (Leray et al., 2021), we wanted to examine the extent to which Ypq1 and Ypq2 can discriminate between lysine and arginine.

We deleted either *YPQ1*, *YPQ2* or both in a *lys2Δarg4Δ* background strain, which ensured that both amino acids cannot be synthesized internally (Beacham et al., 1984; Ljungdahl and Daignan-Fornier, 2012; Sinha and Bhattacharjee, 1971). In each strain, we simultaneously analyzed the renewal dynamics of the CAA in the whole cell extract and separately in the vacuolar and protein fractions. For this, we performed pulse labeling assays under exponential growth conditions, switching cultures from a medium containing normal (light) lysine and arginine to an equivalent one with their heavy stable isotope analogues (Fig. 6 A). We reasoned that, because exponential growth is a steady-state regime, the vacuolar amino acid pools are growing at the same rate as the cell biomass. Therefore, vacuoles must import new amino acids at least as fast as necessary to offset biomass increase. Should lysine and arginine only be imported into the vacuole without any export, the old (light) vacuolar pool would be diluted by new (heavy) amino acids exactly at the rate of cell culture growth (Fig. 6 B). However, if the CAA were also exported from the vacuole, additional import would be required, inevitably leading to higher renewal rates of the vacuolar pool. In other words, vacuolar export in an exponentially growing culture can be detected by jointly analyzing culture growth rate and the dynamics of the old/light amino acid fraction (labeling) in the vacuole (Fig 6 B). In addition, knowing CAA labeling dynamics in the vacuolar, protein, and whole cell lysate fractions all at the same time, would enable to quantitatively compare the size of the vacuolar CAA pools between the mutants and their vacuolar export rates (see below).

**Figure 6.**
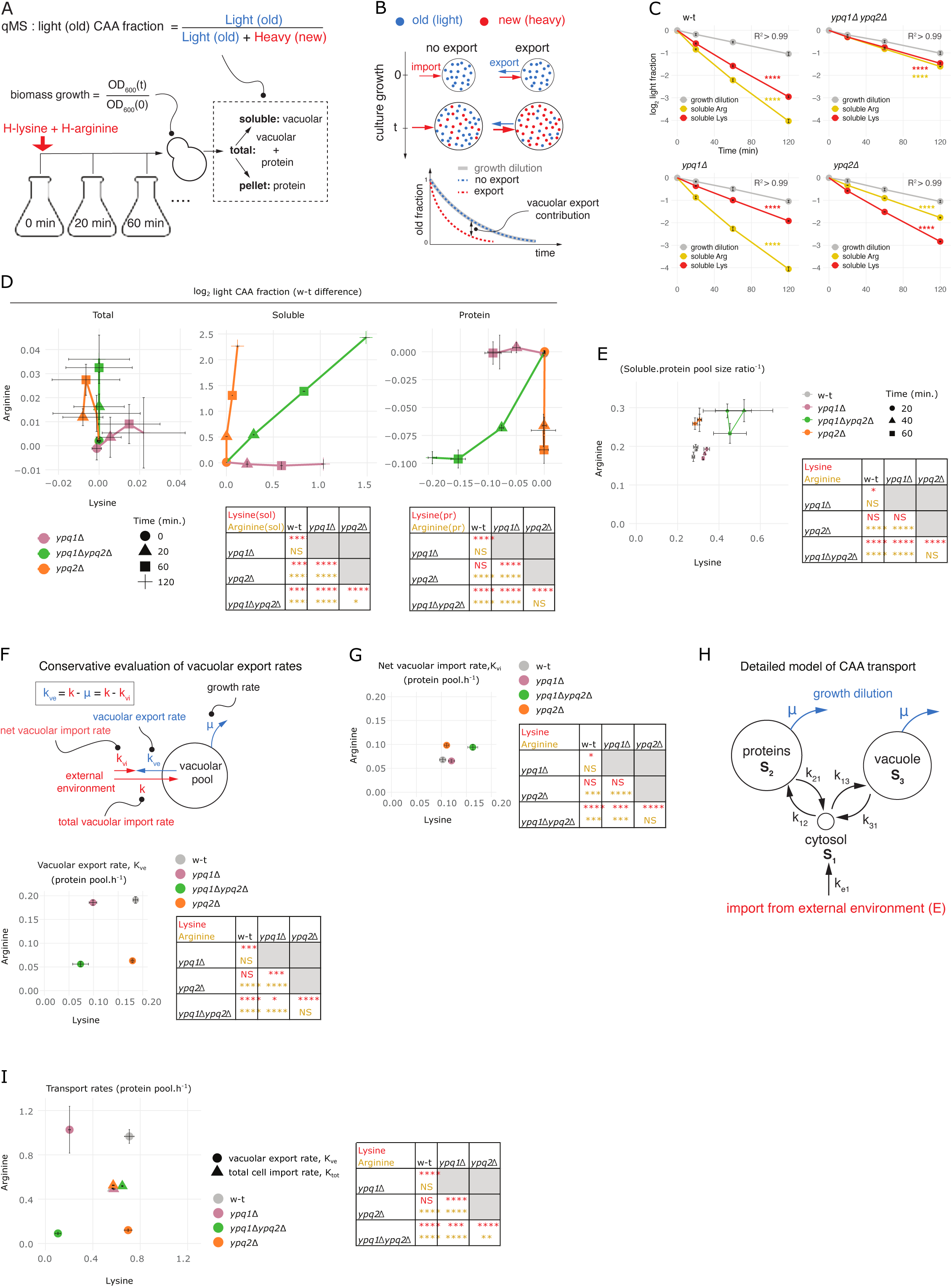
Ypq1 and Ypq2 mediate fast and semi-selective export of vacuolar lysine and arginine in actively growing cells. (A) Schematic of a dynamic labeling assay to monitor CAA renewal. Yeast cultures are grown in SCD medium containing normal (light) lysine and arginine, and in the mid-log phase (OD 600 nm ∼0.15) are transferred by filtration to a pre-warmed “heavy” SCD medium containing [¹³C₆, ¹⁵N₂]-lysine and [¹³C₆, ¹⁵N₄]-arginine at the same concentrations as in the “light” medium. Following medium exchange, the turbidity of samples is determined by OD 600 nm to quantify biomass growth. At each time point after the labeling pulse, the samples are drawn from the growing cultures, cells are washed with distilled water, and the boiled cell suspension (total pool) is split by centrifugation into soluble fraction and pellet representing vacuolar and protein CAA pools, respectively. The fractions are dried and hydrolyzed for 24 h with 6 M HCl at 110 ° C and analyzed by quantitative mass spectrometry to determine the fractional content of the light CAA variants, referred to as labeling. (B) (top) Renewal dynamics of vacuolar CAA pools in exponentially growing cell cultures and its connection to vacuolar export. The pool of “old” light CAA molecules present in the vacuole at the time point of the labeling pulse (time = 0, blue) is diluted by the import of “new” heavy molecules (time = t, red) to just support exponential growth of cell culture (no export). Any export from the vacuole must be additionally compensated for by more import, leading to faster renewal dynamics of the vacuolar CAA pool (export). (bottom) Expected renewal dynamics of vacuolar CAA (measured as the old/light amino acid fraction) corresponding to both scenarios. (C-E) Labeling of soluble, protein, and total pools of CAA compared to the growth dilution determined in dynamic labeling assays with w-t, *ypq1*Δ, *ypq2*Δ and *ypq1*Δ*ypq2*Δ strains. (n = 3). (C) Log-transformed labeling of the soluble pools of CAA at different time points following the labeling pulse plotted together with the log-transformed growth dilution for the same cultures. (****: difference between labeling and growth dilution dynamics p < 0.0001 by two-way ANOVA). Note that the growth dilution curve is essentially linear in all cases (R2 > 0.99). (D) Log-transformed labeling of CAA in different fractions measured in *ypq1*Δ, *ypq2*Δ and *ypq1*Δ*ypq2*Δ strains offset by the corresponding mean values measured in w-t strain. (****: p < 0.0001, ***: p < 0.001, *: p < 0.05 by two-way Tukey HSD). (E) Soluble/protein pool size ratios determined for CAA at different time points using labeling of the total, protein, and soluble fractions as described in (D). (****: p < 0.0001, *: p < 0.05 by two-way Tukey HSD). (F) Conservative evaluation of vacuolar export rates and net vacuolar import rates based on the CAA labeling dynamics, culture growth rates and relative vacuolar CAA content described in (C and E) and expressed in units of the size of the protein pool of CAA. Heavy (new) amino acids considered to be imported into the vacuole directly from the outside environment at a fractional rate k, which is balanced by vacuolar export at a fractional rate k*_ve_* and the net vacuolar import rate k*_vi_*, which is equivalent to the biomass growth at rate μ. (****: p < 0.0001, ***: p < 0.001, *: p < 0.05 by one-way Tukey HSD). (G) Net vacuolar import rates of lysine and arginine determined in units of the size of the cellular protein pool of CAA as described in (F). (****: p < 0.0001, ***: p < 0.001, *: p < 0.05 by one-way Tukey HSD). (H) Detailed model of CAA transport that accounts for the fluxes between protein, vacuolar, and cytosolic pools of CAA. The parameters of the model k*_ij_*are the rates of amino acid transfer from the source pool i to the recipient pool j, measured in units of the size of the recipient pool S*_j_*. All pools are considered to grow at the biomass growth rate μ. (I) Vacuolar export rates of lysine and arginine evaluated using the detailed model (H) based on the labeling dynamics in (C-E) and plotted together with the net cellular import rates of lysine and arginine. (****: p < 0.0001, ***: p < 0.001, **: p < 0.01 by one-way Tukey HSD).

To implement this analysis, we shifted mid-log phase cultures from “light” to “heavy” growth medium differing only in the content of lysine and arginine isotopomers and simultaneously monitored the labeling dynamics of both amino acids in total cell lysate, and in its separate soluble and protein fractions by quantitative mass spectrometry. In parallel, we also recorded culture growth by turbidity measurements (Fig 6 A). Here, the soluble fraction served to represent the vacuolar pools of lysine and arginine, since both are normally confined to the yeast vacuole (Kitamoto et al., 1988; Messenguy et al., 1980) . To ensure that this assumption also held in our mutants, we checked CAA partition between vacuolar and cytosolic fractions by the Cu^2+^ permeabilization, with phenylalanine serving as a fiducial cytosolic marker (Kitamoto et al., 1988; Messenguy et al., 1980). In fact, more than 95 % of both lysine and arginine appeared as vacuolar in all mutants, validating the use of the soluble fraction as a proxy for vacuolar pools (Fig. S 5 A). To verify exponential growth during dynamic labeling assays we plotted the dynamics of the log-transformed biomass growth against labeling pulse duration. In all assays, we observed a highly linear relationship indicating that cells maintain steady-state exponential growth after the labeling pulse (Fig. 6 C).

As judged by the labeling dynamics of soluble CAA, in all four mutants, vacuolar lysine and arginine pools renewed significantly faster than expected from growth dilution alone, indicating that all of them to some extent export lysine and arginine from the vacuole (Fig. 6 C). However, vacuolar lysine pool renewal was markedly slower in the *ypq1Δ* mutant compared to the w-t background and the same was true for arginine in the *ypq2Δ* mutant. Both amino acids renewed slower in *ypq1Δypq2Δ* cells (Fig. 6 D and Fig. S 5 B). Interestingly, these differences were exactly mirrored in the protein fraction but were essentially absent from the total cellular pools (Fig. 6 D and Fig. S 5 B). The lack of detectable effects on the total cellular pools indicates that the deletions of *YPQ1* and *YPQ2* most likely specifically alter the intracellular traffic of CAA. Moreover, slower renewal of vacuolar lysine pool in the *ypq1Δ* strain and of vacuolar arginine pool in the *ypq2Δ* mutant were both consistently accompanied by a faster renewal in the protein pool. We therefore hypothesised that deletions of *YPQ1* and *YPQ2* specifically reduce vacuolar export of lysine and arginine rendering their old vacuolar stores inaccessible for the new protein synthesis.

To investigate this hypothesis further, we quantitatively assessed the vacuolar export rates. These can be conservatively evaluated in units of vacuolar pool size, considering the entire vacuolar import attributed to the heavy CAA acquired directly from the medium after the medium switch. Under this framework, on the one hand, the total valuolar import rates of CAA are equivalent to their renewal rates in vacuolar pool and can be quantified using the observed renewal dynamics (Fig. 6 C). On the other hand, these rates are balanced by the net vacuolar import equivalent to the cell culture growth rate and by the vacuolar export rates (Fig. 6 F). Therefore, fractional vacuolar export rates can be quantified as the difference between the renewal rate of the vacuolar fraction and the growth dilution rate (Fig. 6 F). Since the size of the vacuolar pool can vary between mutants, the fractional export rates must also be adjusted for these differences accordingly. To do this, we determined the vacuolar-to-protein pool size ratios of each amino acid based on the labeling dynamics measured in the soluble, protein, and total cell lysate fractions (Fig. 6 E) and used them to compare export rates in units of the size of the cellular protein pool, which is conceivably similar across mutants. Although the YPQ1 and YPQ2 deletions conferred a significant enlargement of vacuolar lysine and arginine pools, respectively, the export rates appeared to concomitantly drop 2-4 times (Fig. 6 F). We also determined net vacuolar import rates in the same protein pool size units, using the equivalence to the growth rate (Fig. 6 F) and vacuolar-to-protein pool size ratios for their quantification. In fact, net vacuolar import rates appeared to exactly mirror the vacuolar export effects (Fig. 6 G), indicating that mutations do not inhibit vacuolar import capacity of CAA while specifically reducing the vacuolar export capacity. Together, this analysis supports the role of Ypq1 and Ypq2 as relatively specific mediators of vacuolar export of lysine and arginine in actively growing cells.

Since our conservative evaluations do not account, for example, for the reimport of exported amino acids back into the vacuole or import of protein-borne amino acids, the real magnitudes of vacuolar export are likely higher. To determine them more accurately, we developed a detailed model of CAA transport using a compartmental modeling approach (Cobelli et al., 2000; Noor et al., 2025; Onischenko et al., 2020). Our detailed model of CAA transport included vacuolar and protein-borne amino acid pools represented as separate compartments that can acquire amino acids from an external environment and exchange them with each other through a small cytosolic pool (Fig. 6 H). We also assumed that during active growth, cells neither significantly export CAA to the outside environment nor degrade them. These assumptions were validated by analyzing the light CAA content in the heavy CAA medium inoculated with the light CAA labeled yeast cells and by quantifying changes in the total CAA content in the whole culture (cells plus medium) during long-term cell culture growth, respectively (Fig. S 5 C and D). The absence of significant degradation and cellular export allowed us to reduce the number of free parameters in the model and fit them reliably using the experimental CAA labeling dynamics in the protein and vacuolar pools (Fig. S 5 C and D). This more accurate analysis showed that explaining the observed labeling dynamics required vacuolar export rates 4–5 times higher than our conservative estimates (Fig. 6 F and I, Fig. S 5 E and F). For comparison, we also determined the total cell import rates of external lysine and arginine using the labeling dynamics of the total cellular pools of these amino acids, suggesting that during active cell growth CAA are exported from the vacuole at surprisingly high rates comparable to the total rates of their cellular import and exceeding the vacuolar import rates about ten times (Fig. 6 G and I).

In addition to the major effects of the deletions, we noticed that the soluble lysine labeling dynamics was modestly reduced in the *ypq2Δ* mutant as compared to w-t cells (Fig. 6 D). This effect was exacerbated in the *ypq1Δ* background and were mirrored in the protein fraction (Fig. 6 D). A similar effect could also be observed for the soluble arginine pool when comparing the *ypq2Δ* and *ypq1Δypq2Δ* mutants (Fig. 6 D). Furthermore, while soluble pools of lysine and arginine were selectively enlarged in single *ypq1Δ* and *ypq2Δ* mutants, the double deletion had a cumulative effect on the lysine pool size (Fig. 6 E). Consistently, our detailed model showed a statistically significant cumulative impairment of lysine and arginine export in the double mutant compared to the single deletion mutants (Fig. 6 I). These patterns speak for likely incomplete substrate selectivity of Ypq1 and Ypq2.

Together, these results led us to conclude that during active cell growth, Ypq1 and Ypq2 act as fast but likely not completely selective mediators of vacuolar export of lysine and arginine (Fig. 8).

### Ypq1 mediates vacuolar import of lysine in the absence of a proton gradient

Our findings that Ypq1 and Ypq2 confer high export rates of CAA across the vacuolar membrane (Fig. 6 I) imply that it must be compensated by similarly high rates of vacuolar import. Since Vsb1-mediated import being dependent on V-ATPase function is likely coupled to consumption of ATP, this could result in a futile import-export cycle unless there are alternative import routes. Intriguingly, previous studies have shown that Ypq1 can also import lysine into isolated vacuolar vesicles and reconstituted proteoliposomes (Arines et al., 2024; Sekito et al., 2014a). Additionally, Ypq2 has been shown to catalyze bidirectional transport of arginine into isolated vacuoles and vesicles (Cools et al., 2020; Kawano-Kawada et al., 2019). This raises the possibility that Ypq1 could also mediate lysine import into the vacuole independently of the proton gradient.

To test this hypothesis, we took advantage of the fact that the *vph1Δ* mutant, in which the V- ATPase is inactivated (Li and Kane, 2009; Nishi et al., 2003), is unable to establish a detectable intracellular lysine pool (Fig. 1). Thus, we applied the cytochrome C cell permeabilization method to analyze the distribution of ^14^C-lysine between cytosolic and vacuolar fractions of a w- t and *vph1Δ* strain. As in previous experiments, assays were performed in *rsp5* mutant backgrounds, adjusting ^14^C-lysine concentrations accordingly to ensure its comparable uptake (Fig. 7 A). Surprisingly, the *rsp5vph1Δ* mutant accumulated ^14^C-lysine and its derivatives predominantly in the vacuolar fraction, like the *rsp5* mutant (Fig. 7 B). To determine whether this vacuolar accumulation could be attributed to Ypq1, we repeated the assay with an *rsp5vph1Δypq1*Δ double mutant. At comparable levels of uptake (Fig. 7 A), in this case approximately 60 % of ^14^C-lysine and its derivatives were recovered in the cytosolic fraction. Since the *rsp5vph1Δ* mutant is defective in vacuolar acidification (Tarsio et al., 2011) and fails to establish a significant vacuolar lysine pool, Ypq1-mediated uptake of lysine into the vacuole likely occurs independently of the proton gradient.

**Figure 7.**
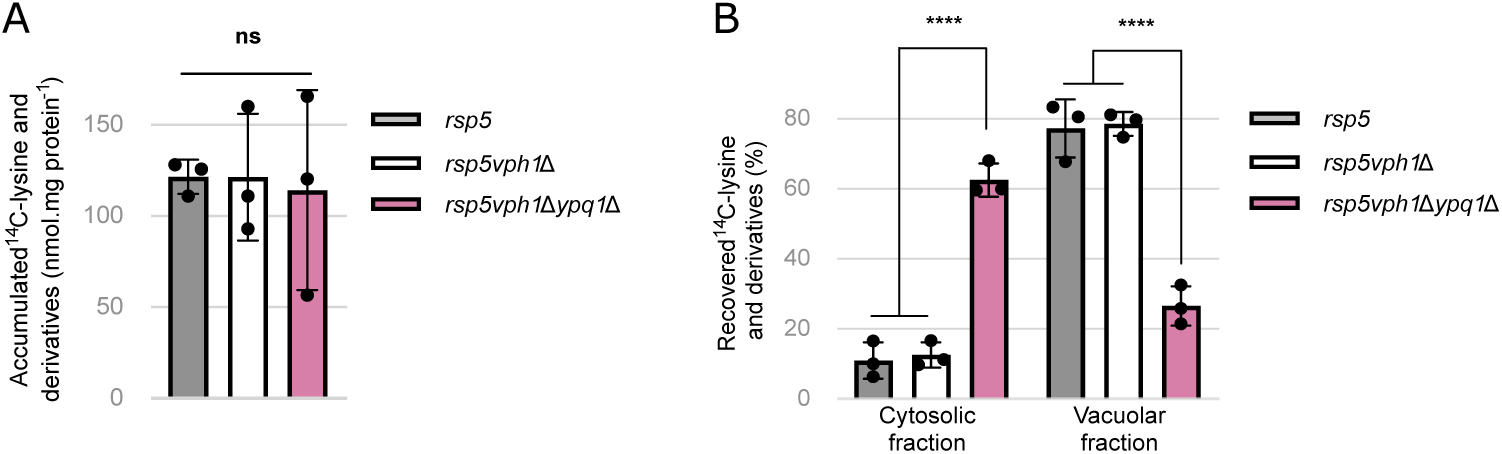
Ypq1 mediates vacuolar import of lysine in the absence of a proton gradient. (A) The *^14^*C-lysine accumulated in the *rsp5* (30 µM *^14^*C-lysine), *rsp5vph1*Δ (90 µM *^14^*C-lysine) and *rsp5vph1*Δ*ypq1*Δ (90 µM *^14^*C-lysine) strains (ns: p > 0.05 by one-way ANOVA test) (n = 3). (B) The distribution of initially accumulated *^14^*C-lysine between the cytosolic and vacuolar fractions after cell permeabilization with cytochrome C in the *rsp5*, *rsp5vph1*Δ and *rsp5vph1*Δ*ypq1*Δ strains (****: p < 0.0001 by one-way ANOVA) (n = 3).

**Figure 8.**
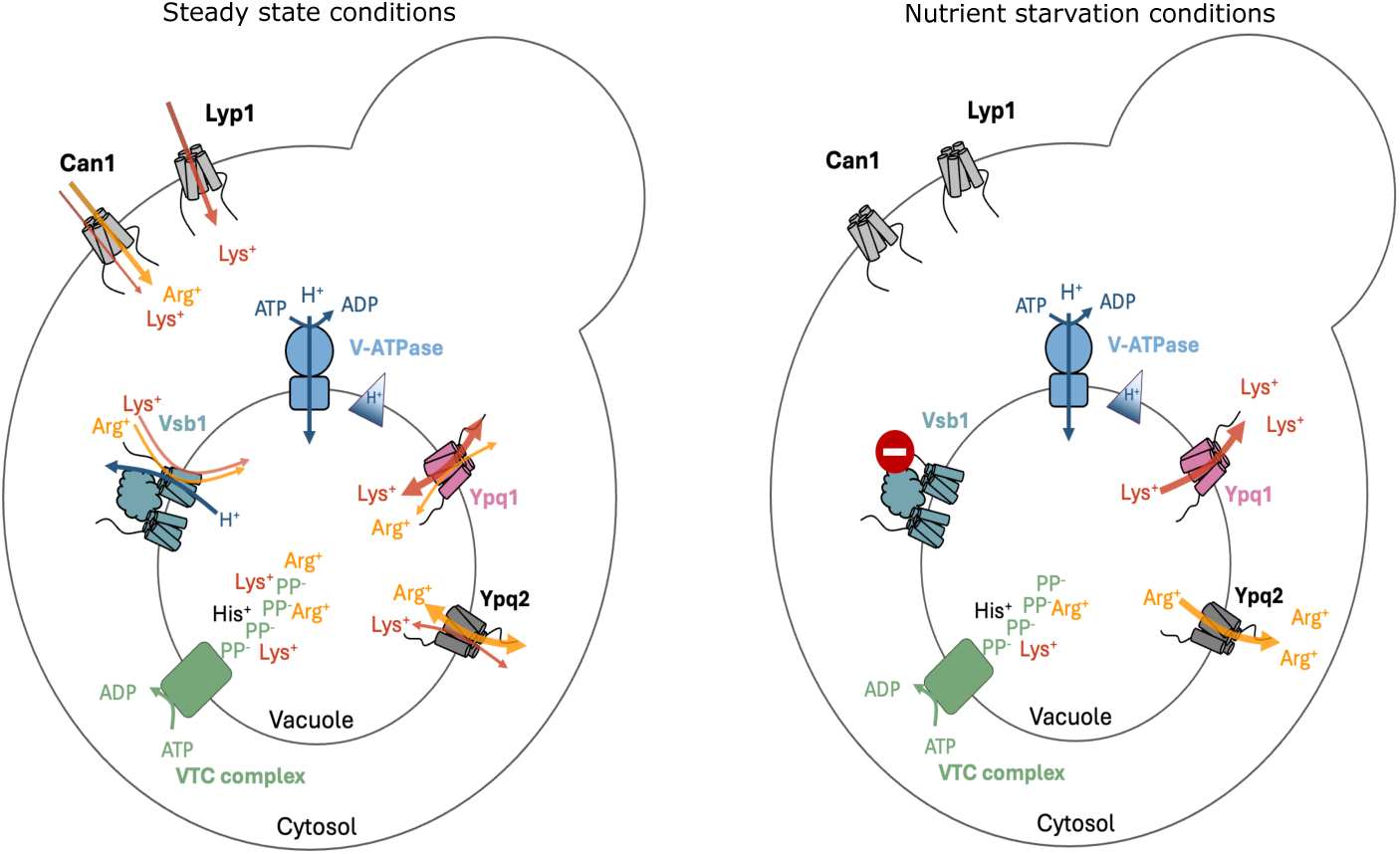
Model for the dynamic storage of cationic amino acid in the vacuole. Under steady state conditions, lysine and arginine are transported into the cytosol from the external environment by the plasma membrane transporters Lyp1 and Can1 and subsequently accumulated in the vacuole by Vsb1, whose activity is likely dependent on the proton gradient established by the V-ATPase. The Ypq1 and Ypq2 facilitators catalyze preferential bidirectional transport of lysine and arginine across the vacuolar membrane, respectively. Lysine and arginine are retained in the vacuole through electrostatic interactions with the negatively charged polyphosphate chains synthesized by the VTC complex. Under lysine starvation, Ypq1 mediates net export of lysine, and under nitrogen starvation, arginine is exported to the cytosol by Ypq2. Vsb1 is presumably inactive under starvation conditions. The export of CAA sustains cell growth under nutrient limitation.

From these observations, we conclude that Ypq1 functions as a facilitator able to catalyze transport of lysine in or out of the vacuole depending on substrate electrochemical gradient.

## Discussion

In this study, we characterized Vsb1 as the main lysine importer at the yeast vacuole. Consistent with prior work (Cools et al., 2020; Kawano-Kawada et al., 2021), Vsb1 localizes to the vacuolar membrane and its loss provokes a reduction of total CAA pools and impairs long-term accumulation of exogenous lysine. Further supporting its role in lysine import, *VSB1* overexpression increases intracellular free lysine levels (Fig. 1). Given that about 90 % of CAA are normally stored in the vacuole (Kitamoto et al., 1988; Messenguy et al., 1980), their total levels serve as a reliable proxy for vacuolar pools. Nevertheless, permeabilization assays showed that, in the *vsb1Δ* mutant, exogenous lysine accumulates in the cytosol instead of the vacuole (Fig. 2 C).

As steady-state lysine levels were only reduced by approximately 50 % in cells lacking Vsb1, additional vacuolar lysine importers are likely to exist. We tested candidates, including the previously described Vba1–3 importers (Shimazu et al., 2005), but deletion of their corresponding genes had no significant effect on lysine levels under our assay conditions (Fig. 1). Although we did not measure Vsb1 activity *in vitro*, previous work using HA-tagged Vsb1 in reconstituted vesicles showed that its arginine transport activity relies on the V-ATPase- generated proton gradient (Kawano-Kawada et al., 2021). Our data are consistent with Vsb1 being a proton antiporter, as the *vph1Δ* mutant, which lacks a functional vacuolar V-ATPase, fails to establish a lysine pool, while bafilomycin A treatment blocks vacuolar accumulation of exogenous lysine in w-t cells (Fig. 1 and 2 C). However, indirect effects of the *VPH1* deletion or bafilomycin A on transporter activity cannot be ruled out.

Since the structure of Vsb1 has not yet been elucidated, we analyzed an Alphafold 3 model of Vsb1 as a dimer, considering the oligomeric state of SLC26A/SulP transporters (Fig. 4 A). Despite its rather poor quality, the model is sound in the light of our experiments showing the importance of the cytoplasmic domains and several key residues in the transmembrane domain. Compared to characterized eukaryotic SLC26A/SulP transporters, Vsb1 has two additional domains, an N-terminal tail of unknown fold and function and a C-terminal RmlC-like domain. While truncating the N-terminal tail did not affect CAA accumulation, the absence of the RmlC-like domain led to a *vsb1*Δ-phenotype. This domain is often associated with nucleotide or metal ion binding (Dunwell et al., 2001), but such a function seems unlikely for Vsb1 since the dimerization does not restore the canonical RmlC binding site (Giraud et al., 2000). Considering the extensive interaction surface of 2180 Å^2^ between the RmlC-like domains in Vsb1, it could have a role in oligomerization like in the potassium channel Eag1 (Whicher and MacKinnon, 2016). Of note, the 385 Å^2^-surface interaction between the STAS domains in the Vsb1 model is unusually small compared to those reported for *Homo sapiens* Slc26A2 (∼ 1100 Å) (Hu et al., 2024), *Meriones unguiculatus* Slc26A5 (∼ 1200 Å) (Butan et al., 2022), and *Mus musculus* Slc26A9 (∼ 1300 Å) (Walter et al., 2019). Even though Vsb1 comprises a core membrane transporter-like domain, we cannot completely exclude the possibility that it does not function as a transporter, as seen in the cochlear outer hair cell motor prestin (Butan et al., 2022) or the yeast Ssy1 receptor of extracellular amino acid (Didion et al., 1998; Iraqui et al., 1999). In that case, Vsb1 may act as a regulator for CAA vacuolar transport. Our mutational study of the putative binding site within the transmembrane domain, however, argues in favor of Vsb1 being a transporter. Indeed, the substitution of Asp-223, Tyr-227 and Glu-278 led to a loss of CAA accumulation and an increased canavanine sensitivity (Fig. 4 E-F). Specifically, Asp-223 is likely an adaptation of the binding site to accommodate a cation instead of an anion, considering that a glutamine residue is found at the same position in all Slc26A anion transporters. As expected from the structures of SLC26A anion transporters, Tyr-227 could be involved in the transport of CAA via H-bond mediated interaction. Since Glu-278 is near the predicted binding site, we hypothesize that its carboxylate could neutralize the TM3 macro-dipole, which is consistent with cation binding. Glu-278 could also play a role in cation exchange. In fact, in BicA, SLC26Dg and AtSULTR4, a carboxylate, brought by either an aspartate or a glutamate residue from the TM8, is found at a position almost equivalent to Glu-278 (Fig. S 2 D) (Geertsma et al., 2015; Wang et al., 2019, 2021) and has been shown to be important for either Na^+^/anion or H^+^ exchange (Wang et al., 2019, 2021).

In addition to elucidating the role of Vsb1 in lysine import, our study provides evidence that the PQ-loop transporter Ypq1 exports lysine out of the vacuole. This conclusion is supported by the elevated lysine pool of the *ypq1Δ* mutant and its strong induction of *LYS9* upon lysine withdrawal (Fig. 5 A). Moreover, under lysine starvation conditions, Ypq1 is critical for the cell to mobilize vacuolar lysine previously accumulated via Vsb1, and to sustain residual growth (Fig. 5 C and Fig. 5 E). Interestingly, the deletion of YPQ1 still allows approximately 30 % of the starting lysine pool to be exported, indicating the involvement of additional exporters. We have ruled out the proposed vacuolar CAA transporters Ypq2, Ypq3 and Avt4 as contributors to lysine export under lysine starvation (Fig. 5 D). This implies that other vacuolar lysine exporters remain to be discovered.

Furthermore, we confirm that Ypq1 undergoes downregulation under lysine starvation, although to a lesser extent than previously reported (Fig. S 4 B) (Arines et al., 2021; Li et al., 2015a, 2015b; Zhu et al., 2017). The mechanism of Ypq1 downregulation has been well characterized, though largely under the assumption that Ypq1 functions as a lysine importer. In this model, Ypq1 transport activity halts when cytosolic lysine levels fall, stabilizing the protein in a conformation that exposes its binding site to the transmembrane adapter Ssh4 (Arines et al., 2021). Ssh4 then recruits the Rsp5 ubiquitin ligase, which ubiquitinates Ypq1, triggering its ESCRT-mediated internalization into the vacuolar lumen for degradation (Li et al., 2015b). Importantly, Ypq1 downregulation in the absence of lysine does not contradict its role in lysine export. In fact, our quantification of cellular lysine pools indicates that complete degradation of Ypq1 occurs several hours after the onset of lysine deprivation, by which time vacuolar lysine stores are nearly depleted (Fig. 5 C) (Li et al., 2015b). We therefore propose that Ypq1, acting predominantly as an exporter, adopts the Ssh4-recognized conformation in response to declining vacuolar, rather than cytosolic, lysine levels. Supporting this model, a recent study demonstrated that purified Ypq1 is destabilized in reconstituted proteoliposomes lacking internal lysine, suggesting a role for luminal lysine in maintaining Ypq1 stability (Arines et al., 2024).

Our dynamic labeling experiments using stable isotope coded amino acids revealed that, not only in starvation conditions but also during active growth, cells continuously export CAA from the vacuole (Fig. 6 C). Kinetic modeling and exploration of the effects of *ypq1Δ* and *ypq2Δ* deletions indicate that Ypq1 and Ypq2 are the key but incompletely selective mediators of this export (Fig. 6). Surprisingly our quantitative analysis of CAA transport based on the dynamic labeling indicates that CAA export rates are very high and comparable to the total cellular import rates of lysine and arginine and exceed the net vacuolar import that supports accumulation of these amino acids in the growing vacuoles almost ten times (Fig. 6 G and I). This raises the intriguing question of how cells maintain even higher vacuolar import to accomplish the CAA accumulation. Should the import of lysine and arginine be solely driven by Vsb1 and other vacuolar proton antiporters, the cell would incur a significant energetic cost just to maintain vacuolar CAA levels. As an alternative, Ypq1 and Ypq2 may function as facilitators (Carruthers, 1990; Leray et al., 2021), catalyzing bi-directional transport of lysine and arginine (Fig. 8) whose net concentration could be retained in the vacuole through electrostatic interactions, for example, with polyphosphate chains (Dürr et al., 1979). Supporting this hypothesis, our current study demonstrates that Ypq1 can mediate lysine import in the absence of a proton gradient across the vacuolar membrane (Fig. 7 B) and we previously observed that Ypq2 can mediate apparent exchange of arginine in isolated vacuoles (Cools et al., 2020). However, a more detailed *in vitro* analysis would be needed to unequivocally determine the transport mechanisms of Ypq1/2 and to understand how environmental conditions and electrochemical gradients modulate their activity.

Finally, our findings illustrate how the vacuole can serve as a dynamic reservoir that regulates CAA homeostasis. In fact, the plurality of CAA transporters at the vacuolar membrane can be explained as a way for a unicellular organism such as yeast to respond quickly to variations in CAA availability. The putative proton antiporter Vsb1 imports lysine and arginine into the vacuole under replete conditions. This transport into the vacuole is key to promote high levels of CAA uptake through the plasma membrane permeases, Lyp1 and Can1 (Fig. 3 B) (Cools et al., 2020; Shimazu et al., 2005). Although energetically costly for the cell as it requires a secondary active transporter, lysine storage mitigates its toxicity, especially under poor nitrogen conditions, and stored CAA pools can be mobilized later. In addition, the Ypq1/2 facilitators are constitutively active in growing cells and dampen fluctuating cytosolic CAA concentrations, preventing excessive biosynthesis under replete conditions (Fig. 1 and 5 A) and promoting quick release to sustain growth under nutrient limitation.

## Materials and methods

### Yeast strains, plasmids, and growth conditions

The yeast strains used in this study (Table. 1) were derived from the reference w-t Σ1278b (Bechet et al., 1970), except for dynamic labeling experiments (see below). All strains were produced using the gene disruption cassette integration method (Guldener, 1996). Except where otherwise stated, cells were grown at 29 °C on a minimal buffered medium (pH = 6.1) (Grenson and Acheroy, 1982) with glucose (3 %) as the carbon source and ammonium in the form of (NH4)2SO4 (10 mM) as the nitrogen source. To complement the uracil auxotrophy, either uracil was added at 0.0025 % or strains were transformed with the pFL38 (Bonneaud et al., 1991) plasmid. When used, lysine was supplemented at a final concentration of 500 µM unless stated otherwise. The final concentrations of substances added to solid or liquid media were for canavanine 0.6 µg/ml and for bafilomycin A 9 µM. In all experiments, cells were examined or harvested during exponential growth, a consistent number of generations after seeding. The plasmids used in this study (Table 2) were constructed using standard molecular cloning techniques.

**Table 1.** Strains used in this study.

| Strain | Genotype | Reference |
| --- | --- | --- |
| Σ1278b derived strains |  |  |
| 23344c | *ura3* | Laboratory collection |
| FV1624 | *ura3vph1Δ* | This study |
| LL162 | *ura3vba1Δvba2Δvba3Δ* | (Cools et al., 2020) |
| COM090 | *ura3vsb1Δ* | (Jézégou et al., 2012) |
| EL029 | *ura3ypq1Δ* | (Jézégou et al., 2012) |
| EL031 | *ura3ypq2Δ* | (Jézégou et al., 2012) |
| EL043 | *ura3ypq3Δ* | (Jézégou et al., 2012) |
| LL180 | *ura3ypq1Δypq2Δypq3Δ* | (Jézégou et al., 2012) |
| FV1505 | *ura3vsb1Δypq1Δ* | This study |
| 27038a | *ura3npi1* | (Hein et al., 1995) |
| SL073 | *ura3npi1car1Δ* | (Cools et al., 2020) |
| SL074 | *ura3npi1car1Δvsb1Δ* | (Cools et al., 2020) |
| COM148 | *ura3lys2Δ* | This study |
| COM146 | *ura3lys2Δypq1Δ* | This study |
| FV1509 | *ura3lys2Δypq1Δypq2Δypq3Δ* | This study |
| FV1456 | *ura3lys2Δvsb1Δ* | This study |
| COM133 | *ura3Ypq1-GFP* | This study |
| COM154 | *ura3lys2ΔYpq1-GFP* | This study |
| FV1626 | *ura3npi1vph1Δ* | This study |
| FV1662 | *ura3npi1vph1Δypq1Δ* | This study |
| CG010 | *ura3gap1Δcan1Δlyp1Δ* | (Gournas et al., 2017) |

| BY4742 derived strains |  |  |
| --- | --- | --- |
| FV1633 | *his3Δ1leu2Δ0lys2Δ0ura3Δ0*<br>*arg4Δ* | This study |
| FV1636 | *his3Δ1leu2Δ0lys2Δ0ura3Δ0*<br>*ypq1Δarg4Δ* | This study |
| FV1638 | *his3Δ1leu2Δ0lys2Δ0ura3Δ0*<br>*ypq2Δarg4Δ* | This study |
| FV1639 | *his3Δ1leu2Δ0lys2Δ0ura3Δ0*<br>*ypq1Δypq2Δarg4Δ* | This study |

**Table 2.**
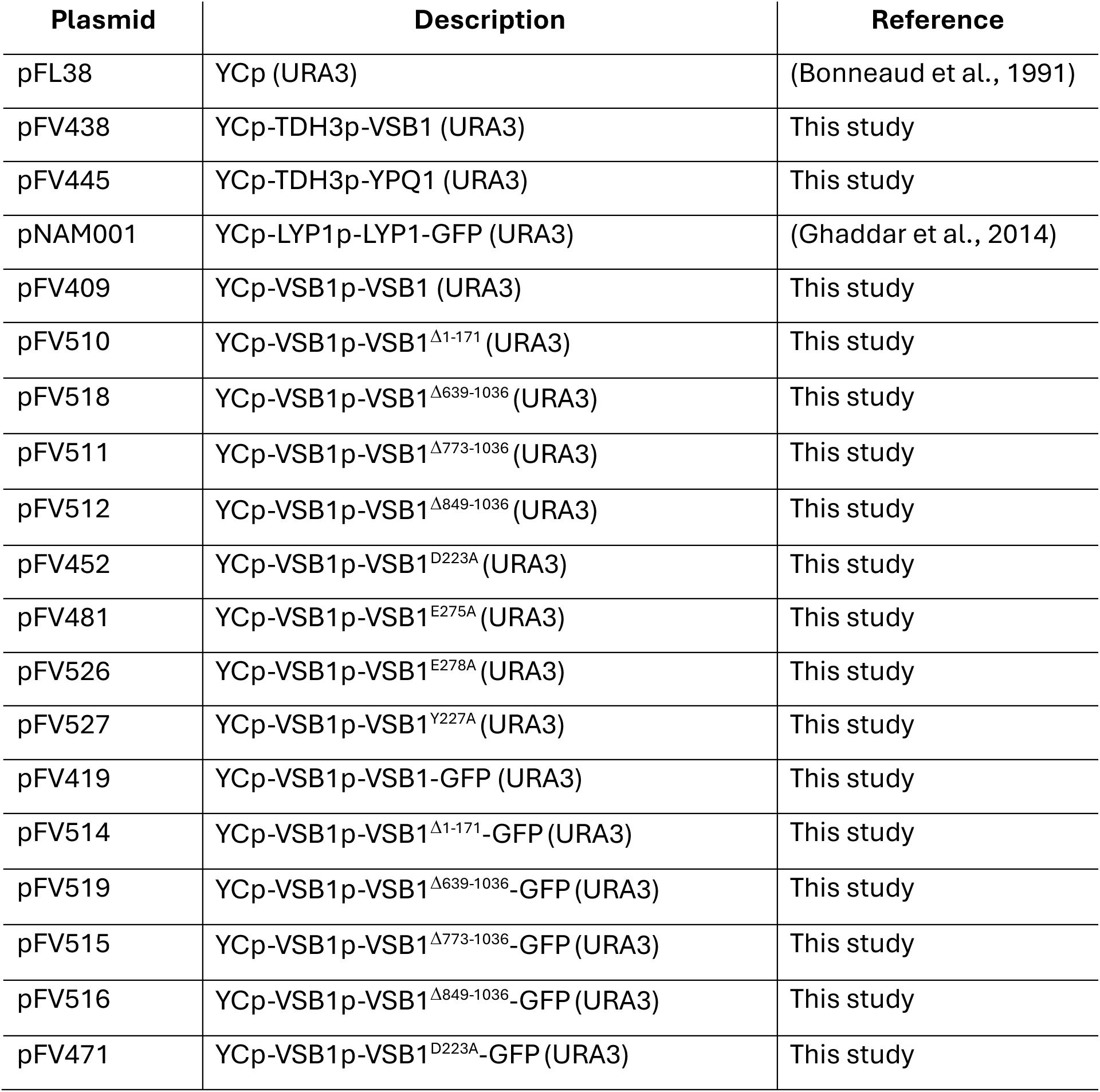
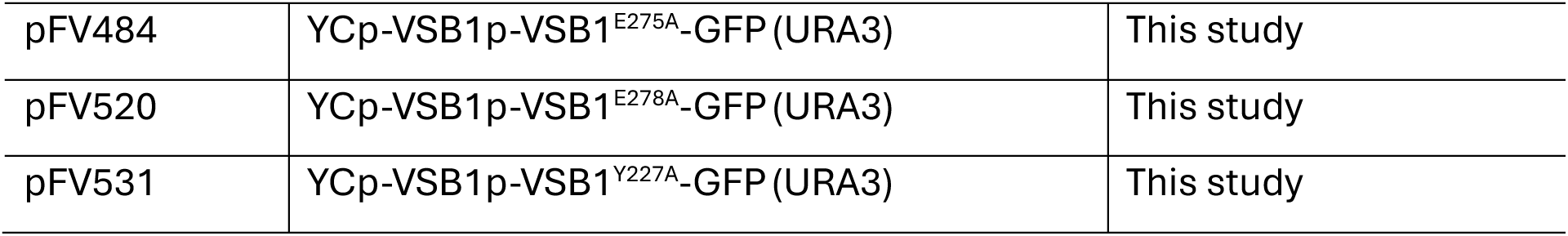
Plasmids used in this study.

### Measurement of total soluble lysine pools

Yeast cultures (25 ml) were collected in exponential phase (∼0.4*10^7^ cells/ml) by centrifugation (7,000 × g for 3 min) and washed three times with 20 ml ultrapure water. Cells were resuspended in 2 ml ultrapure water and boiled for 15 min. The extract was centrifuged (13,000 × g for 3 min) to collect condensation drops and the supernatant was filtered through a PVDF hydrophilic syringe filters (0.2 µm, ROCC) to remove cell debris. Amino acid content was quantified using the AccQ-Tag Ultra turn-key Method (Waters), following the manufacturer’s protocol. Another 25 ml of the same culture was filtrated through an MCE membrane filter (0.45 µm, ROCC) and dried at 60 °C for 24 h to measure dry cell weight for normalization. Data are reported as mean values, with error bars representing standard deviations (SD).

### Uptake assays in whole cells

Accumulation of ^14^C-labeled lysine (Perkin-Elmer) in whole cells was measured at the indicated time points in whole-cell uptake assays, as described previously (Cools et al., 2019; Ghaddar et al., 2014). To determine initial uptake rates, accumulated counts (cpms) were measured 30, 60, and 90 s after the addition of the radioactive substrate. All measurements are expressed in nmol/mg protein per unit of time and reported as mean values, with error bars representing standard deviations (SD).

### Cell permeabilization assays by cytochrome C

Plasma membrane permeabilization by cytochrome C was performed as described previously (Cools et al., 2019). Following a 10 min whole-cell uptake assay, two 5 ml aliquots of culture were filtered onto an MCE membrane filters (0.45 µm, ROCC) and washed with ultrapure water. To measure the total counts per minutes (cpm) of the initial culture, one membrane containing cells was placed in 3 ml of scintillation fluid (Ultima-Flo AP) and radioactivity was measured using a Beckman Coulter LS 6500 Liquid Scintillation Counter (Beckman Coulter). For plasma membrane permeabilization, the second membrane was incubated in 4 ml of cytochrome c solution (1 mg/ml in 1 M sorbitol) for 1 h at 4 °C with gentle shaking. The cell suspension was then percolated over a glass microfiber filter (GF/C 25 mm) and washed three times with 1 ml of 1 M sorbitol. The combined flow-through and wash fractions were collected as the “cytosolic fraction”. To release vacuolar contents, 2 x 3 ml of distilled water was applied to the filter, and the resulting flow-through was collected as the “vacuolar fraction”. For scintillation counting, 500 µL of the cytosolic fraction was mixed with 18 ml of scintillation fluid, and 1 ml of the vacuolar fraction was combined with 6 ml of scintillation fluid. Cpm values of the cytosolic and vacuolar fractions were normalized to the total cpm of the initial culture and reported as mean values, with error bars representing standard deviations (SD).

### Fluorescence microscopy

Cells in exponential phase (∼2*10^6^ cells/ml) were laid down on a thin layer of 1 % agarose. They were viewed at room temperature with an epifluorescence microscope (Eclipse Ci-L; Nikon) equipped with a 100x differential interference contrast N.A. 1.40 Plan Apochromat objective, and appropriate filters. Images were captured with a digital camera (IMAGONGSOURCE TV Lens C-0.45x, Nikon) and NIS-Element D acquisition software (Nikon) and were processed with Fiji software (Schindelin et al., 2012). In each figure, we typically show only a few cells, representative of the whole population. Labeling of the vacuolar lumen with CMAC (7-amino-4- chloromethylcoumarin, ThermoFisher Scientific) was performed by adding the fluorescent dye to a concentration of 25 μM at least 30 min prior to visualization. Labeling of the vacuolar membrane of whole cells with FM4-64 (N-(3-Triethylammoniumpropyl)-4-(6-(4-(Diethylamino) Phenyl) Hexatrienyl) Pyridinium Dibromide, ThermoFisher Scientific) was performed as described previously (Vida and Emr, 1995). For quantifications, images were analyzed with custom-made FIJI macros, calculating the vacuolar membrane-to-total vacuolar intensity. Briefly, for membrane-to-total fluorescence intensity, two homocentric ellipses outlining the whole vacuole or the whole vacuole excluding the vacuolar membrane were manually drawn in middle-section images. The intensities of the channels of interest were measured within manually selected cell outlines, while the median of the fluorescence intensity in the whole image was subtracted as background. All parameters were calculated from at least two independent biological replicates for each condition. The values for single vacuoles are presented in violin plots. After verification that two independent biological replicates gave statistically nonsignificant differences in mean values, the values of the two experiments were merged.

### Growth curves

Comparative analyses of growth in different conditions were performed by growing cells in a 24 or 96-wells non-treated microplate (VWR) incubated at 30 °C with fast shaking (600 rpm) into a SPECTROstar Nano microplate reader (BMG Labtech). Cell growth was monitored by measuring the absorbance at 660 nm every 20 min for 48 h.

### AlphaFold prediction and model analysis

The Vsb1 model was predicted using AlphaFold 3 server (Abramson et al., 2024). Protein structure comparisons were performed using DALI server (Holm et al., 2023). Interaction surfaces were calculated using PDBe PISA (Krissinel and Henrick, 2007). Structures were illustrated using the PyMOL molecular-graphics system version 2.5.0 (Schrödinger).

### Western Blotting

Total cell protein extracts were prepared and analyzed by SDS–PAGE as described previously (Hein et al., 1995). Proteins were transferred to a nitrocellulose membrane (Amersham Protran Premium 0.45 µm) and probed with a mouse monoclonal anti-GFP (RRID: AB_390913, Roche Applied Science), or anti-Pgk1 (PGK1 Monoclonal Antibody 22C5D8, ThermoFisher Scientific). Primary antibodies were detected by enhanced chemiluminescence (SuperSignal West Femto Maximum Sensitivity Substrate, ThermoFisher Scientific, or Immobilon Classico Western HRP Substrate, Merck Millipore) after treatment with horseradish-peroxidase-conjugated anti-mouse or anti-rabbit immunoglobulin (Ig) G secondary antibody (Merck Millipore). Signals were detected with an Imager CHEMI Premium (VWR).

Relative semi-quantitative amounts of total proteins were estimated from 1 to 3 biological replicates of non-saturated exposures using the gel analyzer tool of FIJI. Each band was selected by using rectangular ROI (Region Of Interest) selection and « Gels » analyzer, followed by quantification of the peak area of obtained histograms. Data were acquired as area values. In each graph, the ratios of GFP/Pgk1 normalized to the ratio of experiment which is set as 1 are plotted in a bar chart, normalized to the ratio of EXP which is set as 1.

### Quantitative RT-PCR

RNA isolation and cDNA synthesis were performed with minor modifications to the protocol described by Schmitt *et al*., 1990 (Schmitt et al., 1990). Briefly, total RNA was extracted from 4 ml exponential-phase cultures using the hot acidic phenol method (Schmitt et al., 1990).

Complementary DNA (cDNA) was synthesized from 100 to 500 ng of total RNA using the RevertAid H Minus First Strand cDNA Synthesis Kit (ThermoFisher Scientific) following the manufacturer’s instructions. Purified cDNA was subsequently quantified by quantitative RT-PCR using gene-specific primers (Table 3) and the Power Track SYBR Green Master Mix (ThermoFisher Scientific) on a LightCycler96 system (Roche Applied Science). Standard curves for each primer pair were generated using five successive 10-fold dilutions of genomic DNA. These curves were used to assess PCR efficiency and calculate the relative concentrations of target DNA in all other samples. The specificity of the PCR products was assessed by melting curve analysis. Gene expression levels were normalized to *TBP1* mRNA levels and are presented as mean values. Error bars indicate standard deviations (SD) from replicate measurements.

**Table 3.**
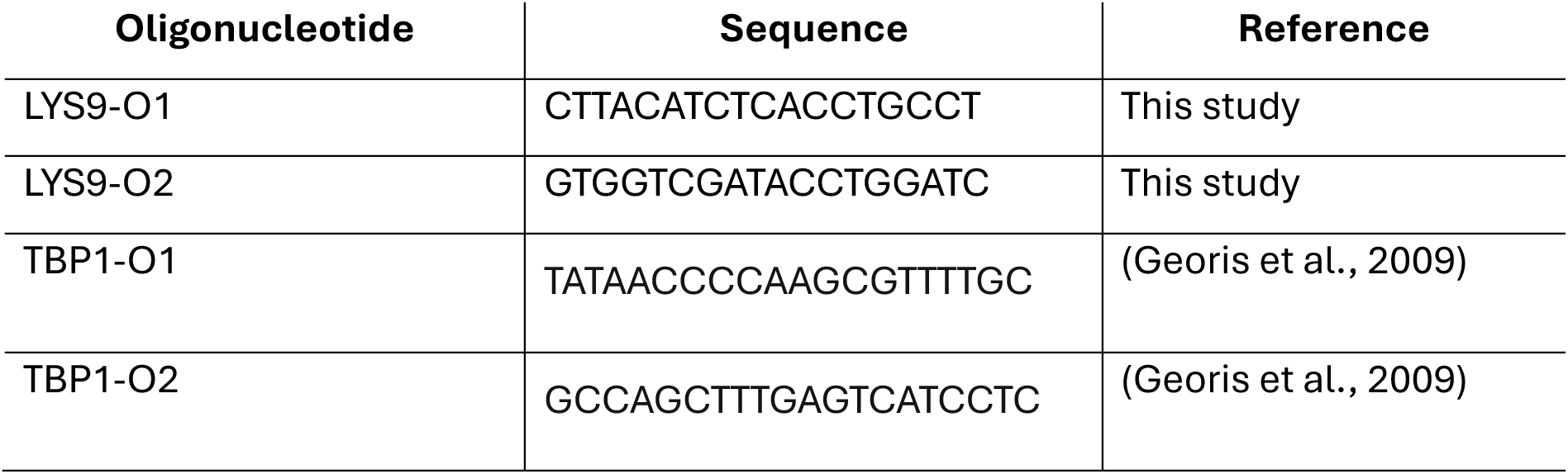
Oligonucleotides used in this study for quantitative RT-PCR.

### Copper chloride permeabilization assay

Yeast strains derived from BY4742 *(MAT(alpha) his3Δ1 leu2Δ0 lys2Δ0 ura3Δ0)* were cultivated at 30 °C in a minimal buffered medium (pH = 6.1) 98 with 3 % glucose as the carbon source and 10 mM (NH4)2SO4 as the nitrogen source. The medium was further supplemented with 0.006 % L-leucine, 0.002 % L-histidine, 0.002 % uracil, 0.0025 % L-lysine, and 0.002 % L-arginine. When cultures reached an absorbance of 0.25 at 660 nm (∼2.5*10^7^ cells/ml), two 12.5 ml aliquots were filtered through an MCE membrane filter (0.45 µm, ROCC) and washed with ultrapure water. The membrane was washed twice with buffer B (2.5 mM potassium phosphate pH = 6, 0.6 M sorbitol), and cells were resuspended in 2 ml of buffer A (10 mM glucose and 0.2 mM CuCl2 in buffer B). After an incubation at 30 °C for 5 min with gentle shaking, the cells were filtered (MCE membrane filters, 0.45 µm, ROCC) and the flowthrough (fraction B) was collected for further analysis. The cells were then resuspended in 2 ml of ultrapure water, and boiled for 15 min. The boiled cell suspension was centrifuged (13,000 × g for 3 min) to collect condensation drops and the supernatant was filtered through a PVDF hydrophilic syringe filter (0.22 µm, ROCC) to remove cell debris. The final extract (fraction C), representing the vacuolar fraction, and cytosolically enriched fraction B were subjected to amino acid analysis by mass spectrometry. The vacuolar lysine and arginine content in the total soluble fraction was determined using phenylalanine as a fiducial cytosolic marker (Kitamoto et al., 1988) and assuming that the exposure to CuCl2 does not significantly release the vacuolar content. In this case, the vacuolar fraction in the total soluble pool, fvac, can be quantified as:

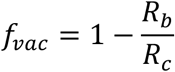

where Rb and Rc are relative amino acid enrichments determined as the signal intensity ratio of the given amino acid and phenylalanine in fractions B and C, respectively. Note that since some vacuoles can be broken by CuCl2 treatment, this provides a conservative evaluation of the size of the vacuolar fraction.

### Metabolic labeling assays

For dynamic labeling assays yeast strains derived from BY4742 *(MAT(alpha) his3Δ1 leu2Δ0 lys2Δ0 ura3Δ0)* were cultivated at 30 °C in a standard synthetic complete medium (pH = 6.1) with yeast nitrogen base containing (NH4)2SO4 as nitrogen source and 2 % glucose as carbon source. The medium was further supplemented with 0.002 % each adenine, L-arginine, L-histidine, L-methionine, L- tryptophan, uracil, 0.006 % L-leucine, 0.0025 % L-lysine, 0.005 % L-phenylalanine, 0.02 % L-threonine, 0.003 % L-tyrosine and designated as the “light medium”. When cultures reached an absorbance of ∼ 0.15 at 600 nm, a 10 ml sample was harvested and stored for further fractionation and amino acid extraction (see below) while 90 ml of the culture was harvested by filtration (WCN membrane filters, 0.8 µm, Whatman), washed on the filter with 10 ml and transferred to 90 ml of a pre-warmed heavy isotope labeled medium containing 0.0025% [¹³C₆, ¹⁵N₂]-L-lysine and 0.002 % [¹³C₆, ¹⁵N₄]-L-arginine instead of light isotopomers. Immediately after transfer, the initial culture turbidly was measured at 600 nm. Culture samples were collected at 20 min, 60 min and 120 min following the medium exchange where 1 ml culture volume was used to determine turbidity and 10 ml was harvested and stored at -20 °C. For harvesting, the culture samples were immediately spun down for 1 min at 2,100 g. The cell pellet was quickly transferred to a 1.5 ml tube, resuspended in 1 ml of MS grade water and spun down again for about 10 s to collect and discard the liquid fraction. The cell pellet was then immediately frozen at -20 °C.

For fractionation and amino acid extraction, the pellets were resuspended in 150 μl of MS grade water each and boiled for 15 min at 100 °C. 100 μl of boiled cell suspension was spun down for 1 min at 17,000 g saving the supernatant, while the pellet was washed once with 100 μl of MS grade water followed by completely removing the liquid fraction. The remaining 50 μl of the boiled cells, and the washed pellet, representing the total and protein fractions, respectively, were dried in a speedvac for two hours at 45 °C, while the supernatant, representing soluble fraction, was stored at -20 °C for further amino acid analysis.

The protein content of the total and protein fractions was hydrolyzed using 6 M hydrochloric acid (Sauer et al., 1996) as follows: The dry material was resuspended in 250 μl of freshly prepared 6 M HCl each, transferred to safe-lock Eppendorf tubes, and incubated for 24 h at 110 °C in a heating block under heavy weight to compensate for overpressure. The liquid phase was then evaporated at 95 °C with constant air flow and the remaining material was resuspended in 150 μl of MS grade water. After pelleting the debris for 2 minutes at 17,000 g, 50 μl of supernatant was saved and stored at -20 °C for amino acid analysis by quantitative mass spectrometry.

The content of the light CAA in each fraction (*labeling*) was determined based on the integrated signal peak intensities of the light (IL) and heavy (IH) isotopomers as:

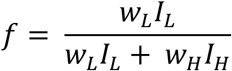

Here, wL and wH are the relative molar weights light and heavy amino acids were determined using a standard equimolar mix of heavy and light amino acids (see mass spectrometry analysis).

For cellular amino acid export tests 100 ml of FV1633 cell culture (*arg4*Δ mutant) grown to OD600 ≈ 0.2 in light SCD medium was harvested, washed with the “heavy” lysine/arginine medium and inoculated in 20 ml of “heavy” lysine/arginine medium exactly as described for dynamic labeling assays. 1 ml cell culture samples collated over 1.5-hour time course were fractionated by centrifugation into soluble (medium) and pellet (cell) fractions. The cell faction of each sample was dried in a speedvac and hydrolyzed with 6 M HCl as described for dynamic labeling, and the extracted amino acids were recovered in 600 μl of MS grade water. Relative amounts of lysine and arginine isotopomers in each fraction (cells and the medium) were determined for all time points using identical 1 μl injection volumes and integrated signal peak intensities of the amino acid signal as readouts.

For amino acid degradation tests 250 μl samples of FV1633 cell culture grown in light SCD medium were periodically collected starting from OD600 ≈ 0.1 over a 20 hour time course. The samples were boiled for 15 min at 100 °C, dried in a speedvac and hydrolyzed with 6 M HCl as described for dynamic labelling assays. After hydrolysis, the amino acid samples were recovered in 250 μl of MS grade water and spiked with an identical volume of heavy lysine /arginine standard mix. The relative total amino acid content in the cell culture samples was then determined as described for the dynamic labeling assays.

### Mass spectrometry analysis

Accurate mass UPLC-HRMS analysis of the samples was performed using a Dionex UltiMate 3000 liquid chromatography system (UPLC) coupled with a Q-Exactive mass spectrometer (HRMS) and interconnected with a heated electrospray ionization source (H-ESI) (Thermo Fisher Scientific, Sunnyvale, CA, USA).

Separation of the analytes was performed on an Acquity Premier BEH Amide VanGuard Fit Column (1.7 µm, 2.1 mm X 100 mm; Waters, Ireland). For analysis, the column was kept at 50 °C, the injection volume was kept at 1 μl and the flow rate was 0.2 ml min^−1^ for a total run time of 15 min. Mobile phase A consisted of 3 % acetonitrile and mobile phase B of 90 % acetonitrile, both buffered with 10 mM ammonium acetate. Separation of individual compounds was achieved using a multistep gradient of A and B where B composition started with 85 % and reduced to 80% in 2 min and further decreased to 50 % over 8 min before being brought to 40 % over 1 min for washout. For equilibration, the B concentration was turned back to 85 % over 4 min.

Ions were monitored in positive targeted single ion monitoring (t-SIM) mode with a resolution of 70,000 at *m*/*z* = 200 and an isolation window of 17 *m*/*z*, using an inclusion parameter list determined using a water-based standard mixture containing 0.5 mM each of L-lysine, heavy L-lysine, L-arginine, heavy L-arginine, L-glutamate, L-histidine, L-threonine, L-tyrosine, L-methionine, L-tryptophan, L-phenylaniline and L-leucine.

Other MS parameters were spray voltage 3.5 kV, sheath gas flow rates 48 units, auxiliary gas flow rate 11 units, sweep gas flow rates 2 units, capillary temperature 256 °C, auxiliary gas heater temperature 413 °C, stacked-ring ion guide (S-lens) radio frequency (RF) level 30, automatic gain control (AGC) 2 × 10^5^ ions, and maximum injection time 200 ms.

Exact mass acquisition and quantification were carried out using the Thermo XCalibur Quan Browser software 4.0.27.42, with 10 ppm mass tolerance.

Before and after each analysis series, six-point dilutions of equimolar [^12^C₆, ^14^N₂] / [¹³C₆, ¹⁵N₂]- L-lysine, and [^12^C₆, ^14^N2] / [¹³C₆, ¹⁵N₄]-L-arginine prepared in MS-grade water were injected and analysed, covering the concentration range from 0.0005 to 0.5 mM and used to determine the relative molar signal weights of the respective isotopomers.

### Determination of transport rates and sizes of amino acid pools

In steady state conditions, vacuolar import of amino acids supports both their continuous export to the cytosol and the growth of the vacuolar mass at the rate of biomass growth (Fig. 6 F), so these rates (expressed in units of the size of the vacuolar pool) can be related as:

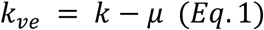

where kve and k are the vacuolar export rate and the bulk vacuolar import rate, respectively, and μ is the biomass growth rate.

For conservative evaluation of vacuolar export rates, all heavy amino acids imported into the vacuole were considered to originate directly from the outside environment. In this case, labeling of the vacuolar pool (the light fraction) after the medium exchange, fsol(t), can be described by a single-exponential decay function as:

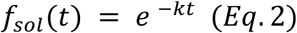

where k is the bulk vacuolar import rate.

By taking the natural logarithm of both sides, Eq.2 can be converted to a linear relationship:

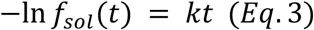

Based on this, bulk vacuolar import rates were evaluated as the slope of linear regression using negative logarithmically transformed labeling values to parametrize the linear model.

The growth rates, μ, were determined analogously considering that the turbidity and the growth rate in exponentially growing cultures are related as:

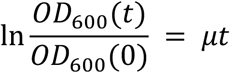

After quantifying the bulk import rates, k, and the growth rates, μ, the vacuolar export rates, kve were determined using Eq.1.

The net vacuolar import rates (expressed in units of the size of the vacuolar pool) kvi were determined directly from biomass growth rate as:

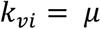

considering that during balanced exponential growth the net vacuolar import supports the net growth of the vacuolar pool at the biomass growth rate (Fig. 6 F).

The net cellular import rates of amino acids ktot were determined in the same way as described for the bulk vacuolar import rates k (see Eq.3), i.e., considering that all heavy amino acids are imported into the cell directly from outside, and that during exponential growth the labeling of the total cellular CAA pool ftot (t) is related to ktot as:

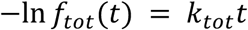

Relative weights of the protein and vacuolar pools of CAA, Sprot and Svac were determined based on the CAA labeling in the total, soluble and protein pool at each time point using ftot(t), fsol(t) and fprot(t), respectively, fractional content relation and considering that most cellular CAA are contained in the protein and vacuolar pools:

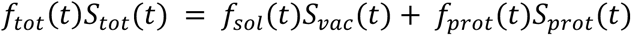

Since the pool weights add up to 1, i.e.

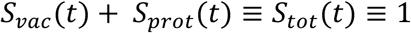

Sprot and Svac can be quantified as:

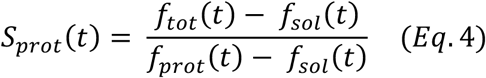

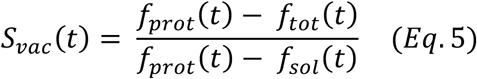

For quantification of transport rates, the pool weights quantified at each time point were averaged to determine their consensus values, Sprot and Svac for each cell culture.

All fractional transport rates were finally converted to units of size of cellular protein pool of respective CAA using the respective pool weights:

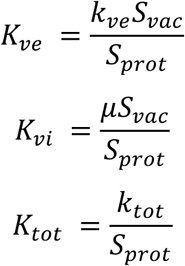

### Construction and parametrization of the detailed model of CAA transport

The detailed model of CAA transport was constructed using a compartmental modelling (CM) approach (Cobelli et al., 2000; Onischenko et al., 2020) and parameterized based on the experimental dynamic labelling readouts using an open-source Symbolic Compartmental Models package available at https://gitlab.com/elad.noor/symbolic-compartmental-model (Noor et al., 2025). In brief, CAA pools, including cytosolic, vacuolar, and protein-borne CAA, were represented as well-mixed compartments that exchange material with each other. In CMs the pools are assigned weights Si which together add up to 1. The transfer of material between pools is described by contributed turnover parameters kij, where the indices i and j represent the source and recipient pool, respectively, and are measured in units of the size of the recipient pool. During steady-state growth, the weights of the pools and the contributed turnovers remain constant.

The graphical representation of our CAA transport model is detailed in the Fig. 6 H. Specifically, the cytosolic pool S1 receives CAA from external environment E at a rate κe1 and exchanges them with the protein pool S2 and the vacuolar pool S3. The contributed turnovers κ13 and κ31 define the rates of vacuolar import and export, respectively, while κ12 and κ21 define the rates of CAA use for protein biosynthesis and their backflux to the cytosolic pool after protein degradation. The blue arrows represent growth dilution of CAA pools.

In steady-state dynamic labeling experiments, the dynamics of the unlabeled fraction in compartments is described through linear inhomogeneous ordinary differential equations that have an analytical solution as the sum of exponential decay functions (Cobelli et al., 2000; Onischenko et al., 2020). Specifically, in the case of our dynamic labeling setup in which unlabeled metabolites are fully replaced with the labeled analogue in growth medium, the dynamics of the unlabeled (light) fraction f (t) in each compartment, which we refer to as *labeling*, can be described through matrix exponentiation in the following form:

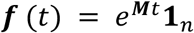

where **f** (t) is a vector of labeling values in each compartment at the time point t after the medium exchange, **1**n is a unit vector and **M** is a transition matrix that describes amino acid transfer rates between compartments.

In our model **f** (t) has 3 components that define the labeling of pools S1, S2 and S3 and the matrix **M** has the following form:

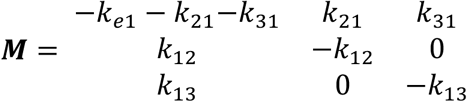

The parameters of the model can be related to a set of measurable and quantifiable parameters by the constraints of mass balance (meaning that the total influx and efflux of each pool must match).

That the influx from external environment supports the growth of the whole system at the cell culture growth rate μ can be expressed as:

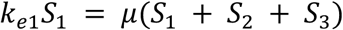

Considering that all pool weights add up to 1, we can express κe1 through this constraint as:

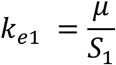

Defining the efflux from the protein pool S2 in units of protein pool size, κpd, as a quantifiable parameter that describes the protein degradation rate and defining κve as the fractional vacuolar export rate and using the above conventions we additionally get:

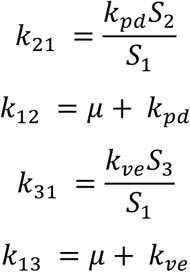

The above mass balance relations were derived assuming that (i) the size of the cytosolic pool of CAA is relatively small compared to the size of the vacuolar and protein CAA pool, i.e. S1 ≪ S2 + S3, and (ii) CAA are not significantly degraded or exported (Fig. S 5 C and D). The weights of the protein and vacuolar pools S2 and S3 were determined as described in section (Determination of transport rates and sizes of amino acid pools) using labeling of CAA in soluble and protein pools at each time point fprot(t) and fsol(t) (Eq. 4 and Eq.5) and were averaged across the time points for each culture. For calculation purposes the weight of the cytosolic pool was set to an arbitrarily small value S1 = 0.01.

The free model parameters included κpd, μ, and κve (which affect the values in the **M**-matrix) and were quantified by fitting the model with experimental labeling values for protein and soluble pools using a nonlinear least-squares solver:

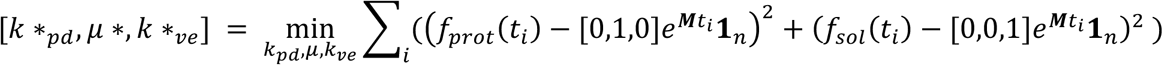

The fitted parameters were constrained to reflect the fact that: (i) vacuolar export rates can have a wide range; (ii) bulk protein degradation rates are low compared to the growth rates of yeast cell cultures under optimal conditions (Wiechecki et al., 2017); and (iii) a reasonable growth rate limit for budding yeast cultures is ∼ 0.45 h^−1^, which corresponds to ∼ 1.5 h doubling time:

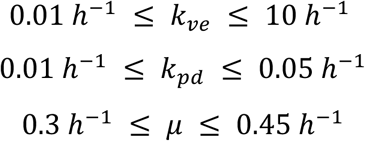

Notably, μ was kept as a free optimization parameter as it could not be determined nearly as accurate the labeling values using turbidity measurements. Nevertheless, to test the validity of our results, we also have set μ to the experimental growth rates derived from turbidity measurements observing the same vacuolar export patterns. The fitting and extraction of the model parameters including the vacuolar export rate and visualization of the fits were performed using a custom Python script. Vacuolar export rates we converted to units of the protein pool size as:

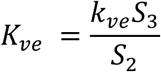

## Supplemental material

Fig. S 1 shows that the *vsb1Δ* mutant is hypersensitive to toxic concentrations of lysine, even when cells are grown on a rich nitrogen source, albeit to a lesser degree. Fig. S 2 shows supplementary analyses of Vsb1 model. Fig. S 3 shows control experiments for Vsb1 model validation. Fig. S 4 shows that Ypq1-GFP is partially targeted to the vacuolar lumen and degraded under lysine starvation. Fig. S 5 shows supplementary analysis of CAA partitioning and transport in w-t yeast cells and in *YPQ1* and *YPQ2* gene deletion mutants.

## Acknowledgments

We thank Christos Gournas for critical reading of the manuscript, regular exchanges and mentorship and members of the Molecular Physiology of the Cell laboratory for fruitful discussions. Evi Zaremba is a fellow of the Fonds pour la formation à la Recherche dans l’Industrie et dans l’Agriculture (FRIA), the Fonds David et Alice Van Buuren, the Fondation Jaumotte-Demoulin and the International Brachet Foundation. This work was supported by the Fonds Stimulus from Meurice R&D and by a research grant from the Research Council of Norway (NFR 315615) to Evgeny Onischenko and Elad Noor.

---

**Figure supplement 1.**
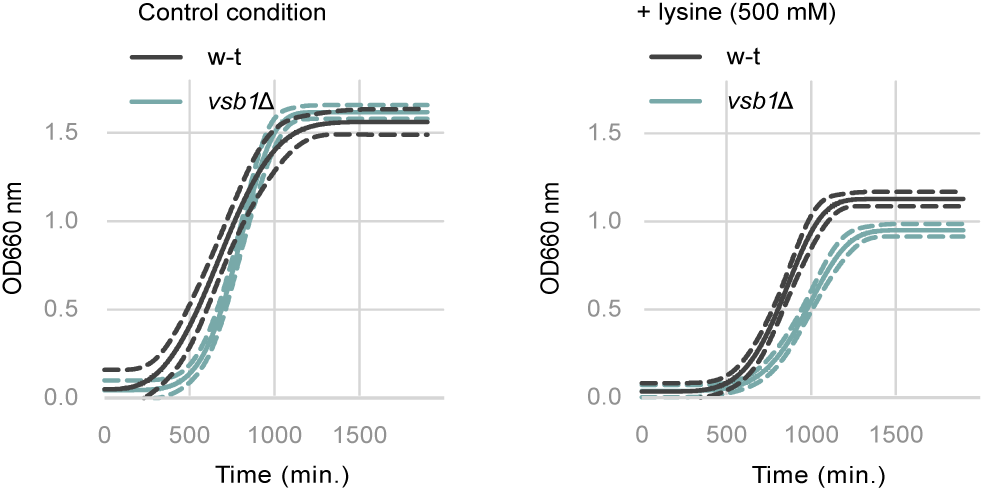
Lysine is toxic at high concentration when grown on a rich nitrogen source. Growth of the w-t and *vsb1*Δ strains in the absence or presence of lysine (500 mM) in the culture media. Cells were initially grown on minimal lysine-free medium containing ammonium as the sole nitrogen source and OD660 nm was monitored over a 24-hour period. Growth is represented as the Weibull nonlinear fit curve with the 99 % confidence bands. (n = 4).

**Figure supplement 2.**
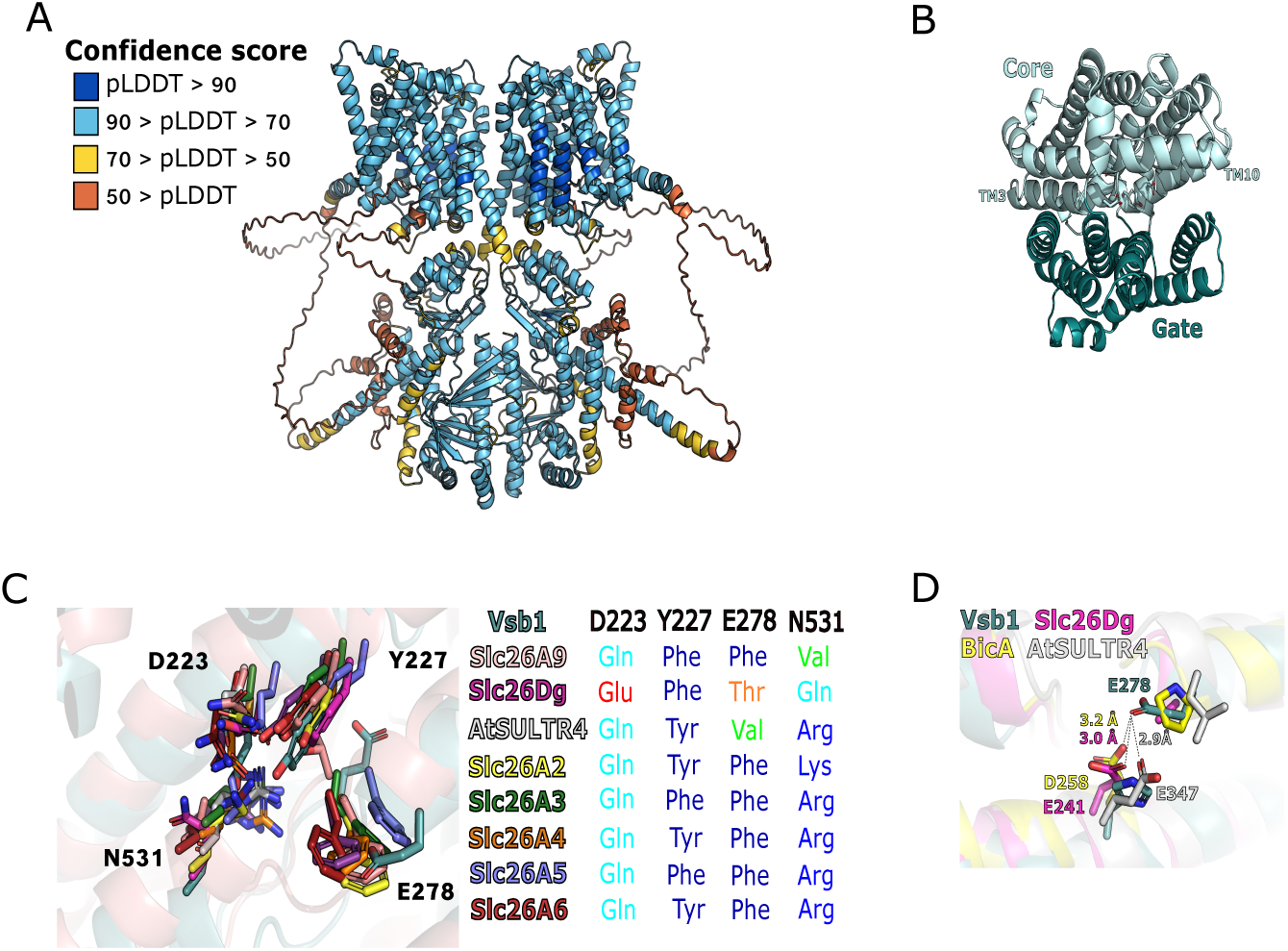
Analysis of Vsb1 structural model and structural comparison with SLC26A/SulP transporters. (A) Vsb1 dimer model predicted using AlphaFold 3 server. The model confidence is represented with colore coded predicted local distance difference tests (orange, pLDDT < 50; yellow, 50 < pLDDT < 70; light blue, 70 < pLDDT < 90; dark blue, pLDDT > 90). (B) Schematic representation of the core (light teal) and gate (dark teal) of Vsb1 TM domain. (C) Close-up view of the putative arginine binding site of Vsb1 (teal) and its structural alignment with human SLC26A9 (pink, PDB code 6RTC, rmsd 3.76 Å), Deinococcus geothermalis SLC26Dg (purple, PDB code 5IOF, rmsd 2.51 Å), Arabidopsis thaliana AtSULTR4 (grey, PDB code 7LHV, rmsd 3.42 Å), human SLC26A2 (yellow, PDB code 7XLM, rmsd 3.82 Å), human SLC26A3 (green, PDB code 8IET, rmsd 4.12 Å), mouse SLC26A4 (orange, PDB code 7WL8, rmsd 3.11 Å), dolphin SLC26A5 (blue, PDB code 7S8X, rmsd 5.11 Å), and human SLC26A6 (red, PDB code 8OPQ, rmsd 3.79 Å). Residues involved in ion binding are shown in stick representation and annotated for Vsb1 only. A table summarizes for each SLC26A transporter which residues are found at the position equivalent to, in Vsb1, Asp-223, Tyr-227, Glu-278, and Asn-531. (D) Close-up view of the binding pocket in-between TM3, TM8 and TM10 and structural alignment of Vsb1 (teal) with Deinococcus geothermalis SLC26Dg (purple, PDB code 5IOF, rmsd 2.51 Å), Arabidopsis thaliana AtSULTR4 (grey, PDB code 7LHV, rmsd 3.42 Å), and Synechocystis elongatus BicA (yellow, PDB code 6KI1, rmsd 4.23 Å). Glu-278 of Vsb1 and the TM8 residues bringing a carboxylate group at an equivalent position in SLC26Dg, AtSULTR4, and BicA (Glu-241, Glu-347, and Asp-258, respectively) are annotated and shown in stick representation.

**Figure supplement 3.**
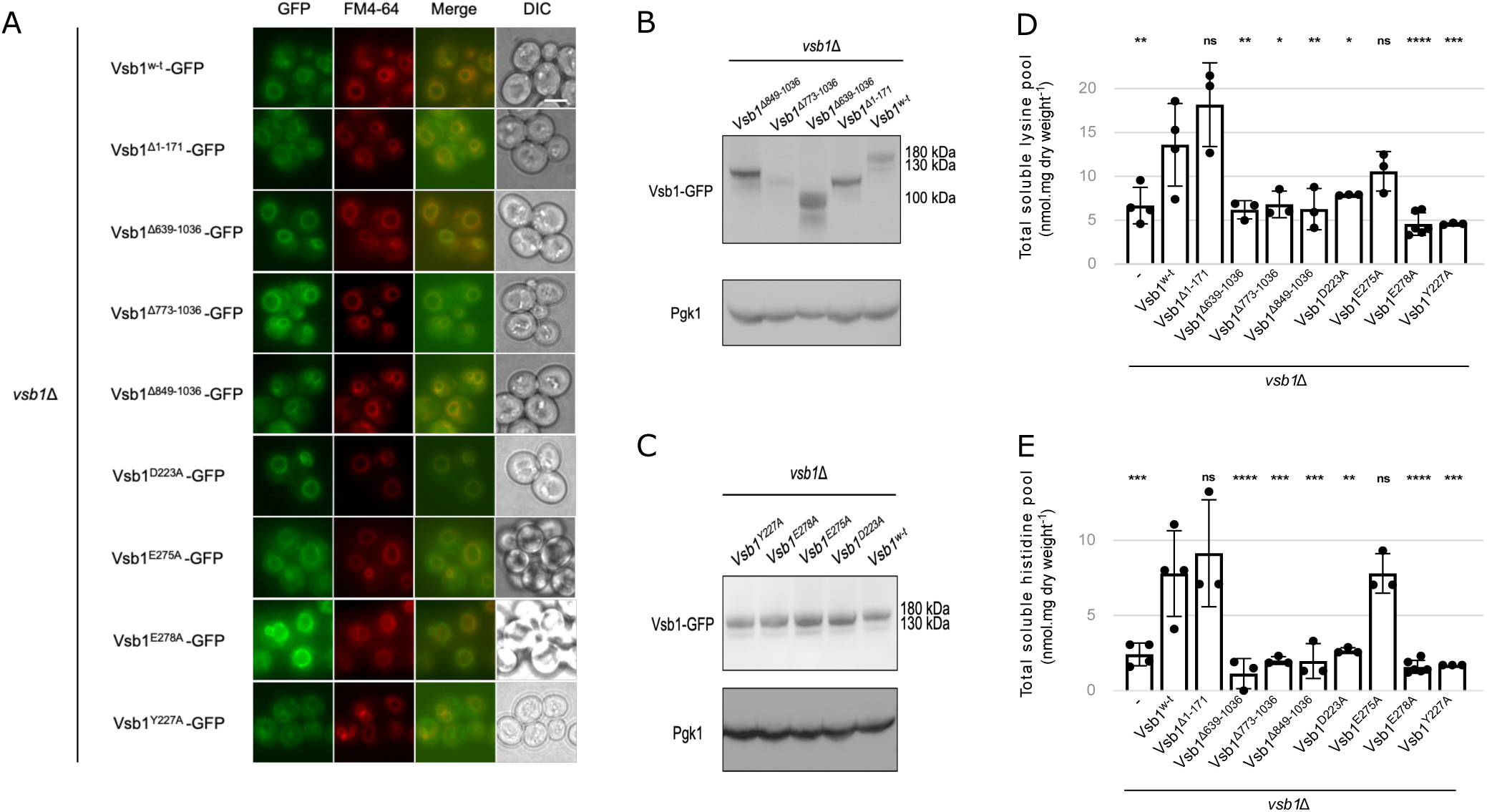
Identification of Vsb1 key residues involved in cationic amino acid transport through mutational analyses. (A to C) A *vsb1*Δ strain was transformed with a plasmid expressing different *VSB1* mutants fused to GFP under the *VSB1* native promoter. (A) Microscopy analysis of different Vsb1-GFP. Scale: 5 µm. (B and C) Western blot of total protein extracts and probed with anti-GFP and anti-Pgk1 (n = 3). (D and E) A *vsb1*Δ strain was transformed with an empty plasmid (-) or plasmid expressing different *VSB1* mutants under the *VSB1* native promoter. Their intracellular lysine (D) and histidine (E) content was measured (ns: p > 0.05; ****: p < 0.0001 by one-way ANOVA with post-hoc comparison tests with the Vsb1^w-t^) (n = 3-4).

**Figure supplement 4.**
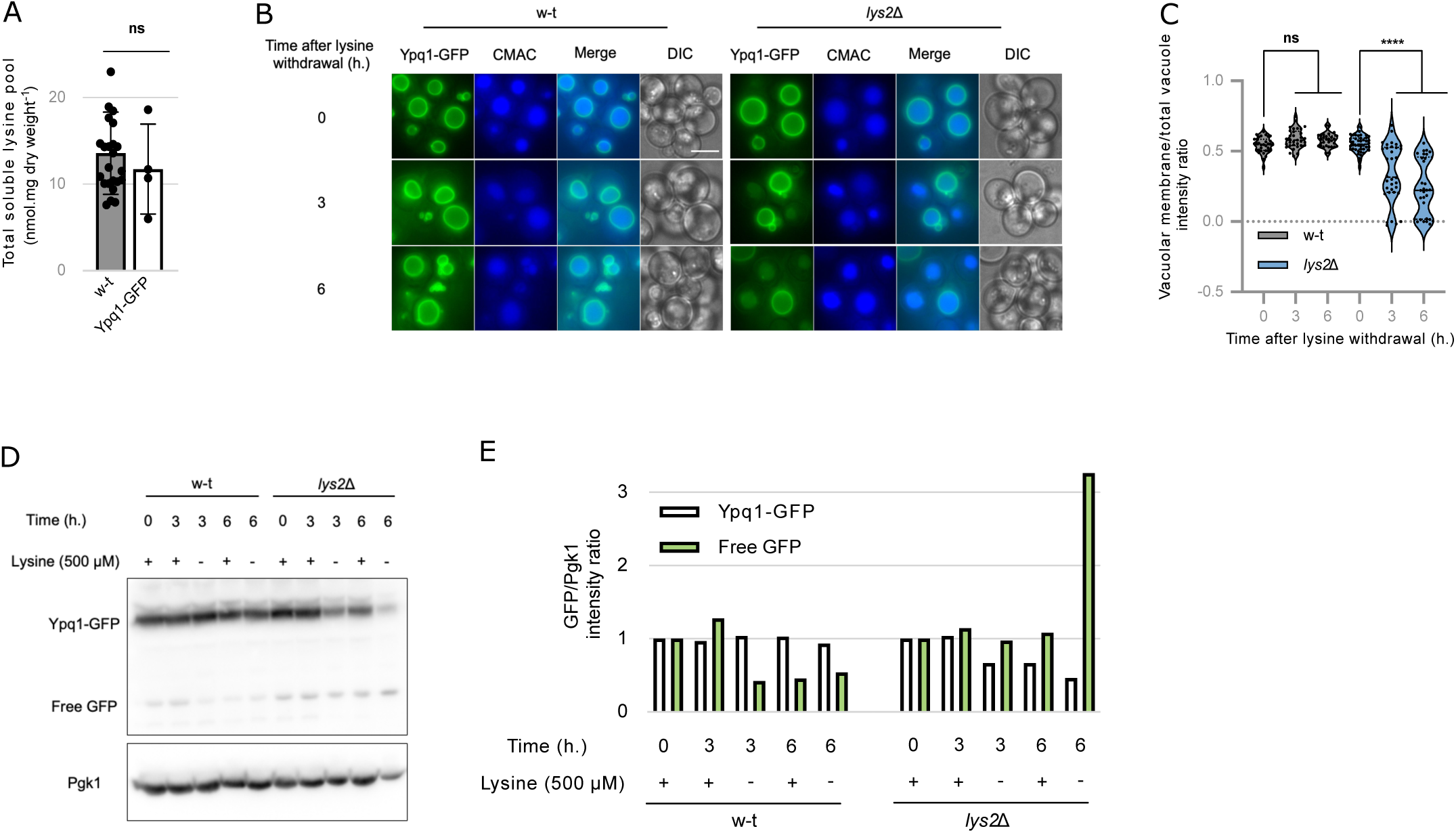
Ypq1-GFP is partially targeted to the vacuolar lumen and degraded under lysine starvation. (A) The intracellular lysine content was measured in the w-t and Ypq1-GFP strains. (ns: p > 0.05 by Student’s test) (n = 4-24). (B) Microscopy analysis of chromosomally tagged Ypq1-GFP localization in the w-t and *lys2*Δ strains stained with vacuolar lumen marker CMAC before and after lysine withdrawal (0, 3 and 6 h). (C) Quantification of (B) Ypq1-GFP vacuolar membrane fluorescence intensity normalized to the total vacuolar fluorescence intensity. (**** : p < 0.0001 by two-way ANOVA test) (n = 54 – 81). (D) Western blot of total protein extracts from a w-t and *lys2*Δ strains expressing chromosomally tagged Ypq1-GFP collected before and after lysine withdrawal (3 and 6 h) and probed with anti-GFP and anti-Pgk1. (E) Quantification of (D) the GFP signal intensity normalized to Pgk1 intensity and the time 0 as experiment 1 (n = 1).

**Figure supplement 5.**
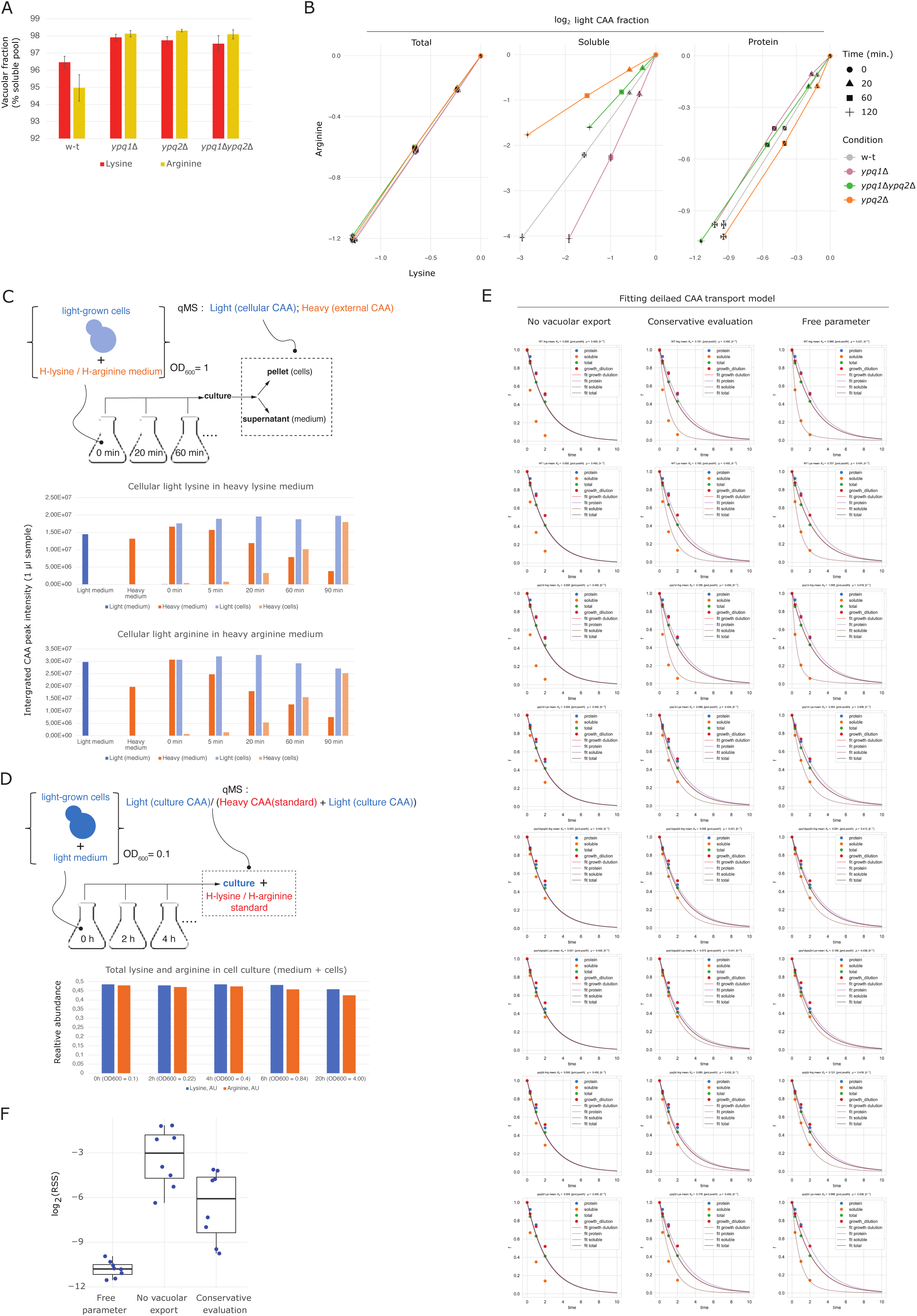
Effect of YQP1 and YPQ2 deletions on cell partitioning and renewal dynamics of CAA. (A) Vacuolar partitioning of CAA determined in actively growing w-t, *ypq1*Δ, *ypq2*Δ and *ypq1*Δ*ypq2*Δ strains with *lys2*Δ*arg4*Δ background using CuCl*_2_* permeabilization. Cells were grown to mid-log phase in presence of 0.0025 % L-lysine and 0.002 % L-arginine, permeabilized for 5 min using CuCl*_2_* in presence of 0.1 M potassium phosphate pH = 6, 4 M sorbitol and 0.018 % glucose. The cytosol-enriched fraction was separated by filtration on MCE membrane filters (0.45 µm, ROCC), while the vacuole-enriched fraction extracted by boiling remaining cell ghosts. The amino acid content of both fractions was quantified by mass spectrometry. (n = 3). Relative enrichments of CAA in CuCl*_2_*permeabilization fractions were determined based on the ratio of signal intensity of the respective CAA and of phenylalanine used as a fiducial cytosolic marker. The relative enrichments were used to conservatively evaluate the fraction of vacuolar CAA in the total soluble amino acid pool (see materials and methods). (B) Log-transformed labeling of CAA in subcellular fractions measured in dynamic labeling assays (see Fig. 6C-E). (C) Export dynamics of cellular CAA to the outside environment. Yeast cells grown in light CAA medium were inoculated into heavy CAA medium followed by measuring the dynamics of light and heavy CAA content in the medium itself (medium) and in the corresponding 6 M HCl hydrolyzed cell pellet (cells) at different time points thereafter. The readout at each time point represents the total signal of CAA (integrated peak intensity) measured by mass spectrometry in 1 µl of the medium supernatant or in the equivalently diluted hydrolyzed cell pellet. Note negligible amounts of light CAA in the medium despite their high intracellular content. (D) Cellular degradation of CAA analysed by monitoring their total content in a growing cell culture using heavy CAA standard. (E) Fitting of detailed model of CAA transport (see Fig. 6H) with the labeling dynamics of CAA in the soluble, protein, and total fraction (see Fig. 6C-E). Three scenarios assume either entire absence of vacuolar export (no export), its conservative estimates (Fig. 6F) as upper limits (conservative evaluation) or setting them as free optimization parameters (free parameter). Note that in the absence of vacuolar export or with its conservative estimates as upper limits, it is not generally possible to explain the experimentally observed CAA labeling dynamics in the soluble and protein pools. (F) Residuals statistics for the three scenarios presented in (E).

## Bibliography

1. Abramson, J., Adler, J., Dunger, J., Evans, R., Green, T., Pritzel, A., Ronneberger, O., Willmore, L., Ballard, A.J., Bambrick, J., Bodenstein, S.W., Evans, D.A., Hung, C.-C., O’Neill, M., Reiman, D., Tunyasuvunakool, K., Wu, Z., Zemgulyté, A., Arvaniti, E., Beattie, C., Bertolli, O., Bridgland, A., Cherepanov, A., Congreve, M., Cowen-Rivers, A.I., Cowie, A., Figurnov, M., Fuchs, F.B., Gladman, H., Jain, R., Khan, Y.A., Low, C.M.R., Perlin, K., Potapenko, A., Savy, S., Singh, S., Stecula, A., Thillaisundaram, A., Tong, C., Yakneen, S., Zhong, E.D., Zielinski, M., Zidek, A., Bapst, V., Kohli, P., Jaderberg, M., Hassabis, D., Jumper, J.M., 2024. Accurate structure prediction of biomolecular interactions with AlphaFold 3. Nature 630, 493–500. 10.1038/s41586-024-07487-w

2. Alper, S.L., Sharma, A.K., 2013. The SLC26 gene family of anion transporters and channels. Mol. Aspects Med. 34, 494–515. 10.1016/j.mam.2012.07.009

3. Arines, F.M., Hamlin, A.J., Yang, X., Liu, Y.-Y.J., Li, M., 2021. A selective transmembrane recognition mechanism by a membrane-anchored ubiquitin ligase adaptor. J. Cell Biol. 220, e202001116. 10.1083/jcb.202001116

4. Arines, F.M., Wielenga, A., Burata, O.E., Garcia, F.N., Stockbridge, R.B., Li, M., 2023. Lysosome transporter purification and reconstitution identifies Ypq1 pH-gated lysine transport and regulation (preprint). Biochemistry. 10.1101/2023.03.31.535002

5. Arines, F.M., Wielenga, A., Stockbridge, R.B., Li, M., 2024. Protocol for purifying and reconstituting a vacuole membrane transporter Ypq1 into proteoliposomes. STAR Protoc. 5, 103483. 10.1016/j.xpro.2024.103483

6. Bavi, N., Clark, M.D., Contreras, G.F., Shen, R., Reddy, B.G., Milewski, W., Perozo, E., 2021. The conformational cycle of prestin underlies outer-hair cell electromotility. Nature 600, 553–558. 10.1038/s41586-021-04152-4

7. Beacham, I., Schweitzer, B., Warrick, H., Carbon, J., 1984. The nucleotide sequence of the yeast ARG4 gene. Gene 271–279.

8. Bechet, J., Grenson, M., Wiame, J.M., 1970. Mutations Affecting the Repressibility of Arginine Biosynthetic Enzymes in *Sacchromyces cerevisiae*. Eur. J. Biochem. 12, 31–39. 10.1111/j.1432-1033.1970.tb00817.x

9. Bertoni, M., Kiefer, F., Biasini, M., Bordoli, L., Schwede, T., 2017. Modeling protein quaternary structure of homo- and hetero-oligomers beyond binary interactions by homology. Sci. Rep. 7, 10480. 10.1038/s41598-017-09654-8

10. Bonneaud, N., Ozier-Kalogeropoulos, O., Li, G., Labouesse, M., Minvielle-Sebastia, L., Lacroute, F., 1991. A family of low and high copy replicative, integrative and single-stranded *S. cerevisiae* / *E. coli* shuttle vectors. Yeast 7, 609–615. 10.1002/yea.320070609

11. Broach, J.R., 2012. Nutritional Control of Growth and Development in Yeast. Genetics 192, 73–105. 10.1534/genetics.111.135731

12. Butan, C., Song, Q., Bai, J.-P., Tan, W.J.T., Navaratnam, D., Santos-Sacchi, J., 2022. Single particle cryo-EM structure of the outer hair cell motor protein prestin. Nat. Commun. 13, 290. 10.1038/s41467-021-27915-z

13. Carruthers, A., 1990. Facilitated diffusion of glucose. Physiol. Rev. 70, 1135–1176. 10.1152/physrev.1990.70.4.1135

14. Cherry, J.M., Hong, E.L., Amundsen, C., Balakrishnan, R., Binkley, G., Chan, E.T., Christie, K.R., Costanzo, M.C., Dwight, S.S., Engel, S.R., Fisk, D.G., Hirschman, J.E., Hitz, B.C., Karra, K., Krieger, C.J., Miyasato, S.R., Nash, R.S., Park, J., Skrzypek, M.S., Simison, M., Weng, S., Wong, E.D., 2012. Saccharomyces Genome Database: the genomics resource of budding yeast. Nucleic Acids Res. 40, D700–D705. 10.1093/nar/gkr1029

15. Chi, X., Jin, X., Chen, Y., Lu, X., Tu, X., Li, X., Zhang, Y., Lei, J., Huang, J., Huang, Z., Zhou, Q., Pan, X., 2020. Structural insights into the gating mechanism of human SLC26A9 mediated by its C-terminal sequence. Cell Discov. 6, 55. 10.1038/s41421-020-00193-7

16. Cobelli, C., Foster, D., Toffolo, G., 2000. Tracer kinetics in biomedical research: from data to model., 1st ed. ed. Kluwer Academic/Plenum Publishers.

17. Cools, M., Lissoir, S., Bodo, E., Ulloa-Calzonzin, J., DeLuna, A., Georis, I., André, B., 2020. Nitrogen coordinated import and export of arginine across the yeast vacuolar membrane. PLoS Genet. 16, e1008966. 10.1371/journal.pgen.1008966

18. Cools, M., Rompf, M., Andre, B., 2019. Measuring the activity of plasma membrane and vacuolar transporters in yeast., in: Yeast Systems Biology. pp. 247–261.

19. Cooper, T.G., Britton, C., Brand, L., Sumrada, R., 1979. Addition of basic amino acids prevents G-1 arrest of nitrogen-starved cultures of Saccharomyces cerevisiae. J. Bacteriol. 137, 1447–1448. 10.1128/jb.137.3.1447-1448.1979

20. Didion, T., Regenberg, B., Kielland-Brandt, M.C., 1998. The permease homologue Ssy1p controls the expression of amino acid and peptide transporter genes in Saccharomyces cerevisiae. Mol. Microbiol. 27, 643–650.

21. Dunwell, J.M., Culham, A., Carter, C.E., Sosa-Aguirre, C.R., Goodenough, P.W., 2001. Evolution of functional diversity in the cupin superfamily. Trends Biochem. Sci. 26, 740–746.

22. Dürr, M., Urech, K., Boller, Th., Wiemken, A., Schwencke, J., Nagy, M., 1979. Sequestration of arginine by polyphosphate in vacuoles of yeast (Saccharomyces cerevisiae). Arch. Microbiol. 121, 169–175. 10.1007/BF00689982

23. Feller, A., Ramos, F., Piérard, A., Dubois, E., 1999. In Saccharomyces cerevisae, feedback inhibition of homocitrate synthase isoenzymes by lysine modulates the activation of LYS gene expression by Lys14p. Eur. J. Biochem. 261, 163–170. 10.1046/j.1432-1327.1999.00262.x

24. Futamata, H., Fukuda, M., Umeda, R., Yamashita, K., Tomita, A., Takahashi, S., Shikakura, T., Hayashi, S., Kusakizako, T., Nishizawa, T., Homma, K., Nureki, O., 2022. Cryo-EM structures of thermostabilized prestin provide mechanistic insights underlying outer hair cell electromotility. Nat. Commun. 13, 6208. 10.1038/s41467-022-34017-x

25. Ge, J., Elferich, J., Dehghani-Ghahnaviyeh, S., Zhao, Z., Meadows, M., Von Gersdorff, H., Tajkhorshid, E., Gouaux, E., 2021. Molecular mechanism of prestin electromotive signal amplification. Cell 184, 4669–4679.e13. 10.1016/j.cell.2021.07.034

26. Geertsma, E.R., Chang, Y.-N., Shaik, F.R., Neldner, Y., Pardon, E., Steyaert, J., Dutzler, R., 2015. Structure of a prokaryotic fumarate transporter reveals the architecture of the SLC26 family. Mol. Biol. 22.

27. Georis, I., Feller, A., Vierendeels, F., Dubois, E., 2009. The Yeast GATA Factor Gat1 Occupies a Central Position in Nitrogen Catabolite Repression-Sensitive Gene Activation. Mol. Cell. Biol. 29, 3803–3815. 10.1128/MCB.00399-09

28. Ghaddar, K., Merhi, A., Saliba, E., Krammer, E.-M., Prévost, M., André, B., 2014. Substrate- Induced Ubiquitylation and Endocytosis of Yeast Amino Acid Permeases. Mol. Cell. Biol. 34, 4447–4463. 10.1128/MCB.00699-14

29. Giraud, M.-F., Leonard, G.A., Berlind, C., Naismith, J.H., 2000. RmlC, the third enzyme of a dTDP- L-rhamnose pathway, is a new class of epimerase. Nat. Struct. Biol. 7, 398–402.

30. Gorbunov, D., Sturlese, M., Nies, F., Kluge, M., Bellanda, M., Battistutta, R., Oliver, D., 2014. Molecular architecture and the structural basis for anion interaction in prestin and SLC26 transporters. Nat. Commun. 5, 3622.

31. Gournas, C., Saliba, E., Krammer, E.-M., Barthelemy, C., Prévost, M., André, B., 2017. Transition of yeast Can1 transporter to the inward-facing state unveils an α-arrestin target sequence promoting its ubiquitylation and endocytosis. Mol. Biol. Cell 28, 2819–2832. 10.1091/mbc.e17-02-0104

32. Grenson, M., 1983. Inactivation-Reactivation Process and Repression of Permease Formation Regulate Several Ammonia-Sensitive Permeases in the Yeast *Saccharomyces cerevisiae*. Eur. J. Biochem. 133, 135–139. 10.1111/j.1432-1033.1983.tb07438.x

33. Grenson, M., 1966. Multiplicity of the amino acid premeases in Saccharomyces cerevisiae II. Evidence for a specific lysine-transporting system. Biochim. Biophys. Acta 127, 339–346.

34. Grenson, M., Acheroy, B., 1982. Mutations affecting the activity and the regulation of the general amino-acid permease of Saccharomyces cerevisiae: Localisation of the Cis-acting dominant pgr regulatory mutation in the structural gene of this permease. Mol. Gen. Genet. MGG 188, 261–265. 10.1007/BF00332685

35. Grenson, M., Mousset, M., Wiame, J.M., Bechet, J., 1966. Multiplicity of the amino acid permeases in Saccharomyces cerevisiae I. Evidence for a specific arginine-transporting system. Biochim Biophys Acta 127, 325–338.

36. Guldener, U., 1996. A new efficient gene disruption cassette for repeated use in budding yeast. Nucleic Acids Res. 24, 2519–2524. 10.1093/nar/24.13.2519

37. Hein, C., Springael, J., Volland, C., Haguenauer-Tsapis, R., André, B., 1995. NPI1, an essential yeast gene involved in induced degradation of Gap1 and Fur4 permeases, encodes the Rsp5 ubiquitin—protein ligase. Mol. Microbiol. 18, 77–87. 10.1111/j.1365-2958.1995.mmi_18010077.x

38. Holm, L., Laiho, A., Törönen, P., Salgado, M., 2023. DALI shines a light on remote homologs: One hundred discoveries. Protein Sci. 32, e4519. 10.1002/pro.4519

39. Horák, J., 1997. Yeast nutrient transporters. Biochim. Biophys. Acta BBA - Rev. Biomembr. 1331, 41–79. 10.1016/S0304-4157(96)00015-9

40. Hothorn, M., Neumann, H., Lenherr, E.D., Wehner, M., Rybin, V., Hassa, P.O., Uttenweiler, A., Reinhardt, M., Schmidt, A., Seiler, J., Ladurner, A.G., Herrmann, C., Scheffzek, K., Mayer, A., 2009. Catalytic Core of a Membrane-Associated Eukaryotic Polyphosphate Polymerase. Science 324, 513–516. 10.1126/science.1168120

41. Hu, W., Song, A., Zheng, H., 2024. Substrate binding plasticity revealed by Cryo-EM structures of SLC26A2. Nat. Commun. 15, 3616. 10.1038/s41467-024-48028-3

42. Iraqui, I., Vissers, S., Bernard, F., Craene, J.-O.D., Boles, E., Urrestarazu, A., Andre, B., 1999. Amino Acid Signaling in Saccharomyces cerevisiae: a Permease- Like Sensor of External Amino Acids and F-Box Protein Grr1p Are Required for Transcriptional Induction of the AGP1 Gene, Which Encodes a Broad-Specificity Amino Acid Permease. MOL CELL BIOL 19, 989–1001.

43. Jauniaux, J.-C., Grenson, M., 1990. GAP1, the general amino acid permease gene of Saccharomyces cerevisiae. Nucleotide sequence, protein similarity with the other bakers yeast amino acid permeases, and nitrogen catabolite repression. Eur. J. Biochem. 190, 39–44. 10.1111/j.1432-1033.1990.tb15542.x

44. Jézégou, A., Llinares, E., Anne, C., Kieffer-Jaquinod, S., O’Regan, S., Aupetit, J., Chabli, A., Sagné, C., Debacker, C., Chadefaux-Vekemans, B., Journet, A., André, B., Gasnier, B., 2012. Heptahelical protein PQLC2 is a lysosomal cationic amino acid exporter underlying the action of cysteamine in cystinosis therapy. Proc. Natl. Acad. Sci. 109. 10.1073/pnas.1211198109

45. Kawano-Kawada, M., Ichimura, H., Ohnishi, S., Yamamoto, Y., Kawasaki, Y., Sekito, T., 2021. Ygr125w/Vsb1-dependent accumulation of basic amino acids into vacuoles of *Saccharomyces cerevisiae*. Biosci. Biotechnol. Biochem. 85, 1157–1164. 10.1093/bbb/zbab015

46. Kawano-Kawada, M., Manabe, K., Ichimura, H., Kimura, T., Harada, Y., Ikeda, K., Tanaka, S., Kakinuma, Y., Sekito, T., 2019. A PQ-loop protein Ypq2 is involved in the exchange of arginine and histidine across the vacuolar membrane of Saccharomyces cerevisiae. Sci. Rep. 9, 15018. 10.1038/s41598-019-51531-z

47. Kim, J., Klionsky, D.J., 2000. Autophagy, Cytoplasm-to-Vacuole Targeting Pathway, and Pexophagy in Yeast and Mammalian Cells. Annu. Rev. Biochem. 69, 303–342. 10.1146/annurev.biochem.69.1.303

48. Kitamoto, K., Yoshizawa, K., Ohsumi, Y., Anraku, Y., 1988. Dynamic aspects of vacuolar and cytosolic amino acid pools of Saccharomyces cerevisiae. J. Bacteriol. 170, 2683–2686. 10.1128/jb.170.6.2683-2686.1988

49. Klionsky, D.J., 1990. The Fungal Vacuole: Composition, Function, and Biogenesis. Microbiol Rev 54.

50. Krissinel, E., Henrick, K., 2007. Inference of Macromolecular Assemblies from Crystalline State. J. Mol. Biol. 372, 774–797. 10.1016/j.jmb.2007.05.022

51. Kschischo, M., Ramos, J., Sychrová, H., 2016. Membrane Transport in Yeast, An Introduction, in: Yeast Membrane Transport, Advances in Experimental Medicine and Biology. Springer International Publishing, Cham, pp. 1–10. 10.1007/978-3-319-25304-6_1

52. Leray, X., Conti, R., Li, Y., Debacker, C., Castelli, F., Fenaille, F., Zdebik, A.A., Pusch, M., Gasnier, B., 2021. Arginine-selective modulation of the lysosomal transporter PQLC2 through a gate-tuning mechanism. Proc. Natl. Acad. Sci. 118, e2025315118. 10.1073/pnas.2025315118

53. Li, F., Vierstra, R.D., 2012. Autophagy: a multifaceted intracellular system for bulk and selective recycling. Trends Plant Sci. 17, 526–537. 10.1016/j.tplants.2012.05.006

54. Li, M., Koshi, T., Emr, S.D., 2015a. Membrane-anchored ubiquitin ligase complex is required for the turnover of lysosomal membrane proteins. J. Cell Biol. 211, 639–652. 10.1083/jcb.201505062

55. Li, M., Rong, Y., Chuang, Y.-S., Peng, D., Emr, S.D., 2015b. Ubiquitin-Dependent Lysosomal Membrane Protein Sorting and Degradation. Mol. Cell 57, 467–478. 10.1016/j.molcel.2014.12.012

56. Li, S.C., Kane, P.M., 2009. The yeast lysosome-like vacuole: Endpoint and crossroads. Biochim. Biophys. Acta BBA - Mol. Cell Res. 1793, 650–663. 10.1016/j.bbamcr.2008.08.003

57. Liu, Q., Zhang, X., Huang, H., Chen, Y., Wang, F., Hao, A., Zhan, W., Mao, Q., Hu, Y., Han, L., Sun, Y., Zhang, M., Liu, Z., Li, G.-L., Zhang, W., Shu, Y., Sun, L., Chen, Z., 2023. Asymmetric pendrin homodimer reveals its molecular mechanism as anion exchanger. Nat. Commun. 14, 3012. 10.1038/s41467-023-38303-0

58. Ljungdahl, P.O., Daignan-Fornier, B., 2012. Regulation of Amino Acid, Nucleotide, and Phosphate Metabolism in *Saccharomyces cerevisiae*. Genetics 190, 885–929. 10.1534/genetics.111.133306

59. Maeda, S., Sugita, C., Sugita, M., Omata, T., 2006. Latent Nitrate Transport Activity of a Novel Sulfate Permease-like Protein of the Cyanobacterium Synechococcus elongatus. J. Biol. Chem. 281, 5869–5876.

60. Manabe, K., Kawano-Kawada, M., Ikeda, K., Sekito, T., Kakinuma, Y., 2016. Ypq3p-dependent histidine uptake by the vacuolar membrane vesicles of *Saccharomyces cerevisiae*. Biosci. Biotechnol. Biochem. 80, 1125–1130. 10.1080/09168451.2016.1141041

61. Manolson, M.F., Proteau, D., Preston, R.A., Stenbit, A., Roberts, B.T., Hoyt, M.A., Preuss, D., Mulholland, J., Botstein, D., Jones, E.W., 1992. The VPH1 gene encodes a 95-kDa integral membrane polypeptide required for in vivo assembly and activity of the yeast vacuolar H(+)-ATPase. J. Biol. Chem. 267, 14294–14303. 10.1016/S0021-9258(19)49711-1

62. McAlister, L., Holland, M.J., 1985. Differential expression of the three yeast glyceraldehyde-3- phosphate dehydrogenase genes. J. Biol. Chem. 260, 15019–15027. 10.1016/S0021-9258(18)95696-6

63. Merhi, A., André, B., 2012. Internal Amino Acids Promote Gap1 Permease Ubiquitylation via TORC1/Npr1/14-3-3-Dependent Control of the Bul Arrestin-Like Adaptors. Mol. Cell. Biol. 32, 4510–4522. 10.1128/MCB.00463-12

64. Messenguy, F., Colin, D., Have, J.-P.T., 1980. Regulation of Compartmentation of Amino Acid Pools in Saccharomyces cerevisiae and Its Effects on Metabolic Control. Eur. J. Biochem. 108, 439–447. 10.1111/j.1432-1033.1980.tb04740.x

65. Nishi, T., Kawasaki-Nishi, S., Forgac, M., 2003. Expression and Function of the Mouse V-ATPase d Subunit Isoforms. J. Biol. Chem. 278, 46396–46402. 10.1074/jbc.M303924200

66. Noor, E., Jefimov, K., Bifulco, E., Onischenko, E., 2025. Age-based approach to characterize the dynamics of cellular processes.

67. Ohnishi, S., Kawano-Kawada, M., Yamamoto, Y., Akiyama, K., Sekito, T., 2022. A vacuolar membrane protein Vsb1p contributes to the vacuolar compartmentalization of basic amino acids in Schizosaccharomyces pombe. Biosci. Biotechnol. Biochem. 86, 763–769.

68. Ohsumi, Y., Anraku, Y., 1981. Active transport of basic amino acids driven by a proton motive force in vacuolar membrane vesicles of Saccharomyces cerevisiae. J. Biol. Chem. 256, 2079–2082. 10.1016/S0021-9258(19)69736-X

69. Ohsumi, Y., Kitamoto, K., Anraku, Y., 1988. Changes induced in the permeability barrier of the yeast plasma membrane by cupric ion. J. Bacteriol. 170, 2676–2682. 10.1128/jb.170.6.2676-2682.1988

70. Olin-Sandoval, V., Yu, J.S.L., Miller-Fleming, L., Alam, M.T., Kamrad, S., Correia-Melo, C., Haas, R., Segal, J., Peña Navarro, D.A., Herrera-Dominguez, L., Méndez-Lucio, O., Vowinckel, J., Mülleder, M., Ralser, M., 2019. Lysine harvesting is an antioxidant strategy and triggers underground polyamine metabolism. Nature 572, 249–253. 10.1038/s41586-019-1442-6

71. Onischenko, E., Noor, E., Fischer, J.S., Gillet, L., Wojtynek, M., Vallotton, P., Weis, K., 2020. Maturation Kinetics of a Multiprotein Complex Revealed by Metabolic Labeling. Cell 183, 1785–1800.e26. 10.1016/j.cell.2020.11.001

72. Preston, R.A., Murphy, R.F., Jones, E.W., 1989. Assay of vacuolar pH in yeast and identification of acidification-defective mutants. Proc. Natl. Acad. Sci. 86, 7027–7031. 10.1073/pnas.86.18.7027

73. Ramos, F., Dubois, E., Piérard, A., 1988. Control of enzyme synthesis in the lysine biosynthetic pathway of *Saccharomyces cerevisiae*: Evidence for a regulatory role of gene *LYS14*. Eur. J. Biochem. 171, 171–176. 10.1111/j.1432-1033.1988.tb13773.x

74. Ramos, F., Verhasselt, P., Feller, A., Peeters, P., Wach, A., Dubois, E., Volckaert, G., 1996. Identification of a gene encoding a homocitrate synthase isoenzyme of Saccharomyces cerevisiae. Yeast 12, 1315–1320. 10.1002/(SICI)1097-0061(199610)12:13<1315::AID-YEA20>3.0.CO;2-Q

75. Reggiori, F., Klionsky, D.J., 2013. Autophagic Processes in Yeast: Mechanism, Machinery and Regulation. Genetics 194, 341–361. 10.1534/genetics.112.149013

76. Russnak, R., Konczal, D., McIntire, S.L., 2001. A Family of Yeast Proteins Mediating Bidirectional Vacuolar Amino Acid Transport. J. Biol. Chem. 276, 23849–23857. 10.1074/jbc.M008028200

77. Sato, T., Ohsumi, Y., Anraku, Y., 1984. Substrate specificities of active transport systems for amino acids in vacuolar-membrane vesicles of Saccharomyces cerevisiae. Evidence of seven independent proton/amino acid antiport systems. J. Biol. Chem. 259, 11505– 11508. 10.1016/S0021-9258(18)90890-2

78. Sauer, U., Hatzimanikatis, V., Hohmann, H.P., Manneberg, M., Van Loon, A.P., Bailey, J.E., 1996. Physiology and metabolic fluxes of wild-type and riboflavin-producing Bacillus subtilis. Appl. Environ. Microbiol. 62, 3687–3696. 10.1128/aem.62.10.3687-3696.1996

79. Schindelin, J., Arganda-Carreras, I., Frise, E., Kaynig, V., Longair, M., Pietzsch, T., Preibisch, S., Rueden, C., Saalfeld, S., Schmid, B., Tinevez, J.-Y., White, D.J., Hartenstein, V., Eliceiri, K., Tomancak, P., Cardona, A., 2012. Fiji: an open-source platform for biological-image analysis. Nat. Methods 9, 676–682. 10.1038/nmeth.2019

80. Schmitt, M.E., Brown, T.A., Trumpower, B.L., 1990. A rapid and simple method for preparation of RNA from Saccharomyces cerevisiae. Nucleic Acids Res. 18, 391–392.

81. Sekito, T., Chardwiriyapreecha, S., Sugimoto, N., Ishimoto, M., Kawano-Kawada, M., Kakinuma, Y., 2014a. Vacuolar transporter Avt4 is involved in excretion of basic amino acids from the vacuoles of *Saccharomyces cerevisiae*. Biosci. Biotechnol. Biochem. 78, 969–975. 10.1080/09168451.2014.910095

82. Sekito, T., Nakamura, K., Manabe, K., Tone, J., Sato, Y., Murao, N., Kawano-Kawada, M., Kakinuma, Y., 2014b. Loss of ATP-dependent lysine uptake in the vacuolar membrane vesicles of *Saccharomyces cerevisiae ypq1* Δ mutant. Biosci. Biotechnol. Biochem. 78, 1199–1202. 10.1080/09168451.2014.918489

83. Sharma, A.K., Rigby, A.C., Alper, S.L., 2011. STAS Domain Structure and Function. Cell. Physiol. Biochem. 28, 407–422. 10.1159/000335104

84. Shimazu, M., Sekito, T., Akiyama, K., Ohsumi, Y., Kakinuma, Y., 2005. A Family of Basic Amino Acid Transporters of the Vacuolar Membrane from Saccharomyces cerevisiae. J. Biol. Chem. 280, 4851–4857. 10.1074/jbc.M412617200

85. Sinha, A.K., Bhattacharjee, J.K., 1971. Lysine Biosynthesis in Saccharomyces 125.

86. Storts, D.R., Bhattacharjee, J.K., 1987. Purification and properties of saccharopine dehydrogenase (glutamate forming) in the Saccharomyces cerevisiae lysine biosynthetic pathway. J. Bacteriol. 169, 416–418. 10.1128/jb.169.1.416-418.1987

87. Sumrada, C., 1976. Basic amino acid inihbition of growth in Saccharomyces cerevisiae. Biochem. Biophys. Res. Commun. 68.

88. Sumrada, R., Cooper, T.G., 1978. Basic Amino Acid Inhibition of Cell Division and Macromolecular Synthesis in Saccharomyces cerevisiae. J. Gen. Microbiol. 108, 45–56. 10.1099/00221287-108-1-45

89. Sun, J., Xu, S., Du, Y., Yu, K., Jiang, Y., Weng, H., Yuan, W., 2022. Accumulation and Enrichment of Trace Elements by Yeast Cells and Their Applications: A Critical Review. Microorganisms 10, 1746. 10.3390/microorganisms10091746

90. Sychrova, H., Matejckova, A., Kotyk, A., 1993. Kinetic properties of yeast lysine permeases coded by genes on multi-copy vectors. FEMS Microbiol. Lett. 113, 57–61. 10.1111/j.1574-6968.1993.tb06487.x

91. Tarsio, M., Zheng, H., Smardon, A.M., Martínez-Muñoz, G.A., Kane, P.M., 2011. Consequences of Loss of Vph1 Protein-containing Vacuolar ATPases (V-ATPases) for Overall Cellular pH Homeostasis. J. Biol. Chem. 286, 28089–28096. 10.1074/jbc.M111.251363

92. Terwilliger, T.C., Liebschner, D., Croll, T.I., Williams, C.J., McCoy, A.J., Poon, B.K., Afonine, P.V., Oeffner, R.D., Richardson, J.S., Read, R.J., Adams, P.D., 2024. AlphaFold predictions are valuable hypotheses and accelerate but do not replace experimental structure determination. Nat. Methods 21, 110–116.

93. Thomas, K.C., Ingledew, W.M., 1994. Lysine inhibition of Saccharomyces cerevisiae: role of repressible L-lysine -aminotransferase. World J. Microbiol. Biotechnol. 10, 572–575.

94. Tippett, D.N., Breen, C., Butler, S.J., Sawicka, M., Dutzler, R., 2023. Structural and functional properties of the transporter SLC26A6 reveal mechanism of coupled anion exchange. eLife 12, RP87178. 10.7554/eLife.87178

95. Vida, T.A., Emr, S.D., 1995. A new vital stain for visualizing vacuolar membrane dynamics and endocytosis in yeast. J. Cell Biol. 128, 779–792. 10.1083/jcb.128.5.779

96. Walter, J.D., Sawicka, M., Dutzler, R., 2019. Cryo-EM structures and functional characterization of murine Slc26a9 reveal mechanism of uncoupled chloride transport. eLife 8, e46986. 10.7554/eLife.46986

97. Wang, C., Sun, B., Zhang, X., Huang, X., Zhang, M., Guo, H., Chen, X., Huang, F., Chen, T., Mi, H., Yu, F., Liu, L.-N., Zhang, P., 2019. Structural mechanism of the active bicarbonate transporter from cyanobacteria. Nat. PLANTS 5, 1184–1193.

98. Wang, L., Chen, K., Zhou, M., 2021. Structure and function of an Arabidopsis thaliana sulfate transporter. Nat. Commun. 12, 4455. 10.1038/s41467-021-24778-2

99. Wang, L., Hoang, A., Gil-Iturbe, E., Laganowsky, A., Quick, M., Zhou, M., 2024. Mechanism of anion exchange and small-molecule inhibition of pendrin. Nat. Commun. 15, 346.

100. Watanabe, T., Ozaki, N., Iwashita, K., Fujii, T., Iefuji, H., 2008. Breeding of wastewater treatment yeasts that accumulate high concentrations of phosphorus. Appl. Microbiol. Biotechnol. 80, 331–338. 10.1007/s00253-008-1529-8

101. Waterhouse, A., Bertoni, M., Bienert, S., Studer, G., Tauriello, G., Gumienny, R., Heer, F.T., de Beer, T.A.P., Rempfer, C., Bordoli, L., Lepore, R., Schwede, T., 2018. SWISS-MODEL: homology modelling of protein structures and complexes. Nucleic Acids Res. 46, W296– W303. 10.1093/nar/gky427

102. Watson, T.G., 1976. Amino-acid Pool Composition of Saccharomyces cerevisiae as a Function of Growth Rate and Amino-acid Nitrogen Source. J. Gen. Microbiol. 96, 263–268. 10.1099/00221287-96-2-263

103. Whicher, J.R., MacKinnon, R., 2016. Structure of the voltage-gated K+ channel Eag1 reveals an alternative voltage sensing mechanism. Science 353, 664–669.

104. Wiechecki, K., Manohar, S., Silva, G., Tchourine, K., Samson, J., Valleriani, A., Vogel, C., 2017. Integrative meta-analysis reveals that most yeast proteins are very stable.

105. Wiemken, A., Nurse, P., 1973. Isolation and characterization of the amino-acid pools located within the cytoplasm and vacuoles of Candida utilis. Planta 109, 293–306. 10.1007/BF00387098

106. Yang, Z., Huang, J., Geng, J., Nair, U., Klionsky, D.J., 2006. Atg22 Recycles Amino Acids to Link the Degradative and Recycling Functions of Autophagy. Mol. Biol. Cell 17, 5094–5104. 10.1091/mbc.e06-06-0479

107. Zhu, L., Jorgensen, J.R., Li, M., Chuang, Y.-S., Emr, S.D., 2017. ESCRTs function directly on the lysosome membrane to downregulate ubiquitinated lysosomal membrane proteins. eLife 6, e26403. 10.7554/eLife.26403

